# Local-to-global signal transduction at the core of the Mn^2+^ sensing riboswitch

**DOI:** 10.1101/553354

**Authors:** Krishna C. Suddala, Ian R. Price, Michal Janeček, Petra Kührová, Shiba Dandpat, Jiří Šponer, Pavel Banáš, Ailong Ke, Nils G. Walter

**Affiliations:** Single Molecule Analysis Group and Center for RNA Biomedicine, Department of Chemistry, University of Michigan, Ann Arbor, MI, 48105; Department of Molecular Biology and Genetics, Cornell University, Ithaca, NY 14850; Institute of Biophysics of the Czech Academy of Sciences, Kralovopolská 135, 612 65 Brno, Czech Republic; Regional Centre of Advanced Technologies and Materials, Department of Physical Chemistry, Faculty of Science, Palacký University, tř. 17 listopadu 12, 771 46, Olomouc, Czech Republic

## Abstract

The widespread manganese-ion sensing *yybP-ykoY* riboswitch controls the expression of bacterial Mn^2+^ homeostasis genes. Here, we first determine the crystal structure of the ligand-bound *yybP-ykoY* riboswitch from *Xanthomonas oryzae* at 2.85 Å resolution, revealing two conformations with docked four-way junction (4WJ) and incompletely coordinated metal ions. In >50 μs of MD simulations, we observe that loss of divalents from the core triggers local structural perturbations in the adjacent docking interface, laying the foundation for signal transduction to the regulatory switch helix. Using single-molecule FRET, we unveil a previously unobserved extended 4WJ conformation that samples transient docked states in the presence of Mg^2+^. Only upon adding sub-millimolar Mn^2+^, however, can the 4WJ dock stably, a feature lost upon mutation of an adenosine contacting Mn^2+^ in the core. These observations illuminate how subtly differing ligand preferences of competing metal ions become amplified by the coupling of local with global RNA dynamics.

## INTRODUCTION

Riboswitches are structured RNA motifs commonly found in the 5’-untranslated regions of bacterial mRNAs, where they regulate many essential and virulence genes in response to binding of a specific ligand^1–3^. Currently, there are over 40 different riboswitch classes known to respond to ligands ranging from metabolites^4^, enzyme cofactors^5^, signaling molecules^6–8^, tRNAs^9^, and to metal ions^10–12^. Ligand binding generally stabilizes a conformation of the riboswitch that modulates either Rho-independent transcriptional termination or translation initiation through accessibility of the Shine-Dalgarno (SD) sequence. The static ligand-bound structures, and the ligand-recognition modes, of a number of riboswitches have been determined at atomic resolution^2,12–14^. Often, the ligand occupies a linchpin position in the global fold where distal residues of the RNA are brought together; however, the dynamic paths by which the local binding of a ligand as small as a metal ion are transduced into the large-scale molecular rearrangements necessary for a regulatory decision by the gene expression machinery largely remain enigmatic^14^.

The *yybP-ykoY* RNA motif is one of the most widespread riboswitches across bacteria, including human and plant pathogens^15–17^. It has evolved to sensitively detect Mn^2+^ metal ions and broadly regulate a variety of genes, particularly those involved in Mn^2+^ homeostasis, at the levels of either transcription or translation^12,15,18^. We previously solved crystal structures of this riboswitch, suggesting that the charge, geometry, and Lewis-acid hardness of Mn^2+^ may be sensed locally by encapsulation of the cation by five phosphoryl oxygens and the N7 of a conserved adenosine. The global structure suggested that formation of the binding site requires “docking” of two distal helical legs of a four-way junction (4WJ) to form a paperclip-shaped global architecture, facilitated by an A-minor interaction and a second, nonspecific divalent metal binding site (**Fig. 1a**)^12^. However, the transduction of ligand binding in this metal-sensing core into global structural changes that affect the distal helix P1.1 involved in riboswitching is not understood, rendering it an archetypical representative of our level of understanding of many crystallized riboswitches^14^.

**Figure 1.**
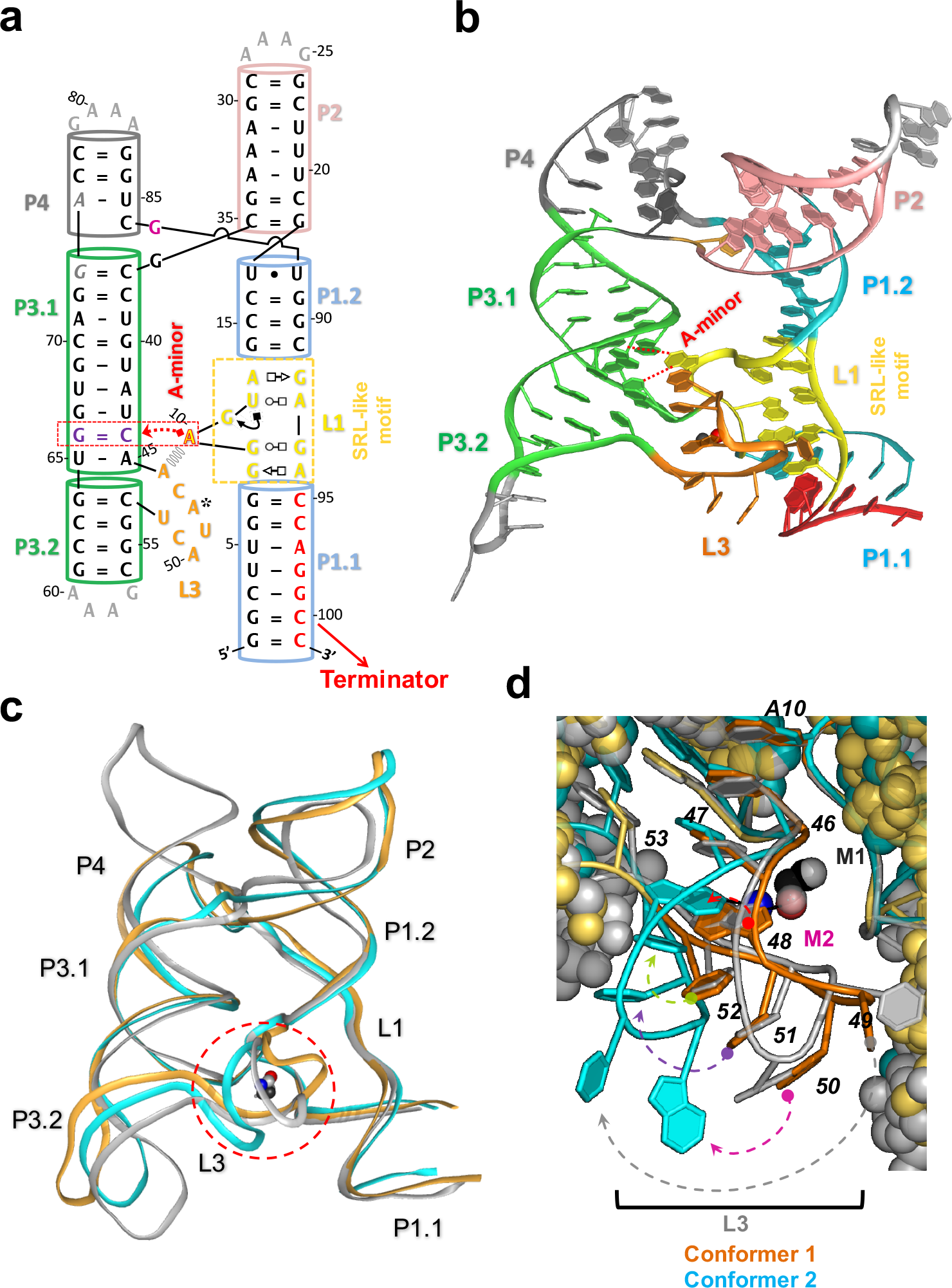
Sequence and crystal structure of the X. oryzae (*Xory*) *yybP-ykoY* Mn^2+^ riboswitch. (**a**) Sequence and secondary structure of the *X. oryzae yybP-ykoY* RNA crystal construct. A native CA dinucleotide was omitted between G73-A74 (gray italics) for crystallization purposes only. The A-minor interaction between L1 and L3 and the SRL-like conformation of L1 are highlighted. (**b**) Crystal structure of Conformer 1 showing the global tertiary fold of the riboswitch along with different secondary structures. (**c**) Comparison of overall structures of Conformers 1 (orange) and 2 (cyan). The two molecules in the asymmetric unit are overall fairly similar. However, they differ dramatically at the metal binding sites (dotted circle). The *L. lactis* structure (gray) is overlaid for comparison. (**d**) Conformer 1 (orange) is relatively similar to the *L. lactis* structure at the M_B,Mn_ binding site. All the same metal contacts are made, although U49, away from the Mn^2+^ site, is shifted (gray dotted arrow). Conformer 2 (cyan) differs in that a metal is still bound at the M_B,Mn_ site, but only half of the metal contacts are made, and the binding site A48 is flipped (red dotted arrow) to expose N1 rather than N7. U49 (gray arrow), A50 (magenta arrow), C51 (purple arrow), and U52 (green arrow) are all significantly shifted from the previously-reported Mn^2+^-bound conformation.

Here, we first solve the structures of two ligand-bound states of the *yybP-ykoY* transcriptional riboswitch from the rice pathogen *X. oryzae*^19,20^, captured in distinct conformations that offer ‘snapshots’ of structural changes *en route* to full ligand binding. We then use these conformers for atomistic molecular dynamics (MD) simulations that reveal how ligand-dependent local structural perturbations in the metal-sensing core are linked to the stability of distal P1.1 helix to affect riboswitching. Finally, using single-molecule FRET (smFRET), we investigate the global structural dynamics of the riboswitch in the presence of varying concentrations of Mg^2+^ and Mn^2+^, as well as other transition metals, revealing a previously unobserved extended conformation of the riboswitch. We show that addition of Mg^2+^ induces two kinetically distinct docked conformations that remain in dynamic equilibrium with the extended conformation. In contrast, upon addition of sub-millimolar Mn^2+^, the riboswitch adopts a stably docked conformation that becomes abolished upon mutation of the conserved core adenosine. Taken together, our work reveals the ligand-dependent (un)folding pathway of the Mn^2+^ sensing riboswitch as a guide for how subtle binding preferences distinguishing two similar metal ion ligands cascade through the coupling of local with global RNA conformational dynamics into powerful effects on gene regulation.

## RESULTS

### *X. oryzae yybP-ykoY* crystal structures reveal disordered metal sensing cores

We determined the structure of the *yybP-ykoY* riboswitch from *X. oryzae* in the presence of Mn^2+^ (Xory-Mn) by X-ray crystallography at 2.85 Å resolution (**Fig. 1b** **and** **Supplementary Table 1**). In this phytopathogenic bacterium that causes rice blight, the riboswitch is found upstream of the *yebN* Mn^2+^ efflux pump gene^20^, whose characterization helped identify the *yybP-ykoY* RNA as a Mn^2+^ riboswitch. To facilitate crystallization, the wild-type (WT) Xory sequence was modified distally from the metal sensing core not to affect metal binding, tightening the 4WJ and stabilizing P1.1 to aid crystallization (see Methods, **Fig. 1a**).

**Table 1.** Complete list of our 29 MD simulations of distinct start conditions and lengths, comprising a cumulative 53 μs simulation time

| Organism | PDB ID | Structure | Chain | Ions in active site |  | solvent box | Length |
| --- | --- | --- | --- | --- | --- | --- | --- |
| | | | | $\text{M}_{\text{A,Mg}}$ | $\text{M}_{\text{B,Mn}}$ | | |
| *L. lactis* | 4Y1I | complete | A | $\text{Mg}^{2+}$ | $\text{Mn}^{2+}$ | rectangular | Each<br>3 $\mu\text{s}$ |
| | | | | $\text{Mg}^{2+}$ | $\text{K}^+$ | | |
| | | | | $\text{K}^+$ | $\text{Mn}^{2+}$ | | |
| | | | | $\text{K}^+$ | $\text{K}^+$ | | |
| | | P1.2\|L1\|P1.1 | A | - | - | rectangular | 3 $\mu\text{s}$ |
| *X. oryzae* | This study | complete | Conformer 2 with *anti* A48 | $\text{Mg}^{2+}$ | $\text{Mn}^{2+}$ | octahedral | Each<br>2 $\mu\text{s}$ |
| | | | | $\text{Mg}^{2+}$ | $\text{K}^+$ | | |
| | | | | $\text{K}^+$ | $\text{Mn}^{2+}$ | | |
| | | | | $\text{K}^+$ | $\text{K}^+$ | | |
| | | | Conformer 2 with *syn* A48 | $\text{Mg}^{2+}$ | $\text{Mn}^{2+}$ | | |
| | | | | $\text{Mg}^{2+}$ | $\text{K}^+$ | | |
| | | | | $\text{K}^+$ | $\text{Mn}^{2+}$ | | |
| | | | | $\text{K}^+$ | $\text{K}^+$ | | |
| | | P1.2\|L1\|P1.1 | Conformer 2 | - | - | rectangular | 3 $\mu\text{s}$ |
| | | complete | Conformer 1 | $\text{Mg}^{2+}$ | $\text{Mn}^{2+}$ | octahedral | Each<br>twice<br>2 $\mu\text{s}$ |
| | | | | $\text{Mg}^{2+}$ | $\text{K}^+$ | | |
| | | | | $\text{K}^+$ | $\text{Mn}^{2+}$ | | |
| | | | | $\text{K}^+$ | $\text{K}^+$ | | |
| | | P1.2\|L1\|P1.1 | Conformer 1 | - | - | rectangular | 3 $\mu\text{s}$ |

Fortuitously, the resulting Xory-Mn structure contained two different conformations in the same crystal lattice (**Fig. 1c**). The two conformers, termed “1” and “2”, are globally quite similar to each other (overall RMSD = 2.53 Å) and, as expected, to the previously determined *L. lactis* canonical Mn^2+^-bound structure, featuring a dual coaxial stack connected at the top by the 4WJ^12^ (**Figs. 1b,c**). As in Llac-Mn, Xory-Mn shows that the conserved L1 and L3 loops dock the two adjacent legs of the 4WJ^12^, wherein L1 forms a sarcin-ricin-like motif (SRL-like) that flips out A10, which forms a cross-helix A-minor interaction with helix P3.1 above L3 (**Fig. 1c**). This non-standard sarcin-ricin loop differs from the canonical loop since A10 is inserted upstream of the G-bulge region (i.e., G9) and the *trans*-Watson-Crick/Hoogsteen base pairing of G9-G93 is distinct from the canonical *trans*-Hoogsteen/Hoogsteen pairing^21^. L3 stacks below A10, orienting its phosphates inward, resulting in a pocket of high negative charge density. Consequently, between the loops two largely dehydrated metal sites are found. In Llac-Mn, the first site, M_A_, is inner-sphere coordinated (at a distance of 2.0-2.3 Å) by five phosphoryl oxygens and a water molecule, and was predicted to be a nonspecific divalent binding site, generally harboring Mg^2+^; we will henceforth refer to it as M_A,Mg_. The second site, M_B_, is coordinated by five phosphoryl oxygens and the N7 of a highly conserved adenosine A41 in L3, which – through its “soft” coordination – appears to help specify Mn^2+^ as the preferred metal ion ligand^12^ (**Fig. 1d**); we will henceforth refer to it as M_B,Mn_. The two new structures each contain both of these ions, and thus support our previous analysis^12^. Of note, we used 80 mM SrCl_2_ and 20 mM MgCl_2_ in the crystallization and, based on the anomalous signal obtained near the Sr K-edge wavelength (0.769 Å), there is no or little Sr^2+^ occupancy at either of the M_A,Mg_ or M_B,Mn_ sites compared to external sites in the structure, attesting to specificity for the smaller divalent ions.

While the two new Xory-Mn conformers are similar, the regions immediately surrounding the metal ion binding sites differ between them and Llac-Mn. The Conformer 1’ M_B,Mn_ binding area is most similar to the *L. lactis* M_B,Mn_, except U49 in the L3 backbone is flipped out, away from the stacked conformation observed for the equivalent *L. lactis* A42 (**Fig. 1d**, **Supplementary Fig. 2**), while still preserving all M_B,Mn_ metal contacts (**Fig. 2a**). By comparison, the conformer 1 M_A,Mg_ site is more distinct in that the inner-sphere contacts to the phosphoryl oxygens of G8 and A10 are lost (shown as red lines), with G8 potentially retaining an outer-sphere, through-water contact (**Fig. 2a**).

**Figure 2.**
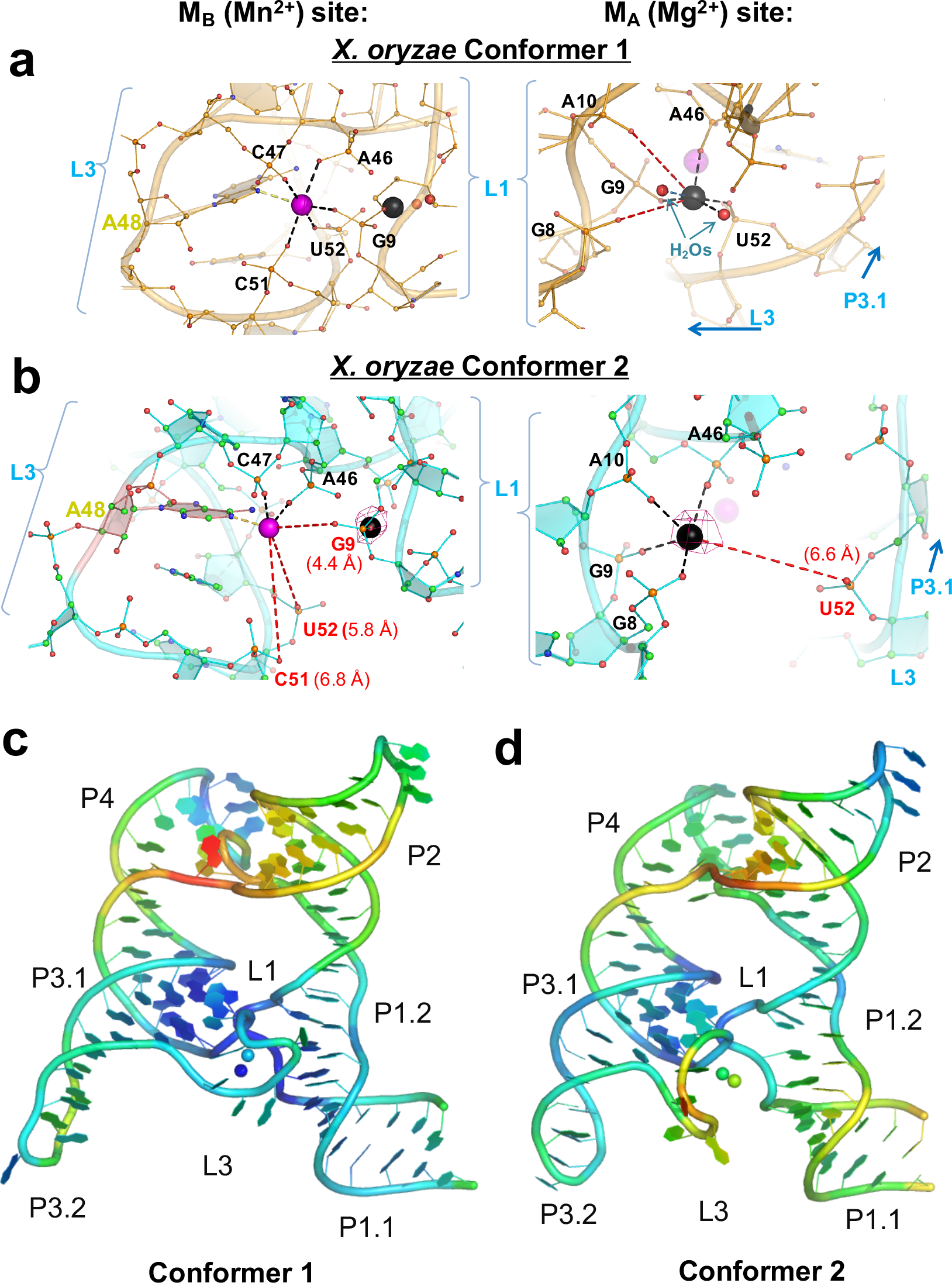
Conformational differences in the metal binding sites and B-factors in the structures. Close-up view of the M_A,Mg_ (black) and M_B,Mn_ (purple) metal binding sites of (**a**) Conformer 1 and (**b**) Conformer 2 showing different contacts with surrounding L1 and L3 residues. Waters are shown as red spheres. The lost metal contacts are shown as red dashed lines with their distances. The anomalous difference map at 4 σ (magenta mesh) shows that Sr^2+^ is not observed in Conformer 1 and weakly occupies the M_A_ site but not the M_B_ site in Conformer 2. Overall *X. oryzae* Conformer 1 (**c**) and Conformer 2 (**d**) structures, colored by individual atomic temperature B-factors (range 38-135 Å^2^ for Conformer 1, 44-170 Å^2^ for Conformer 2). Notice both are well-structured in L1 and in P3 around the binding site. However, Conformer 1 is much more tightly structured in L3 than is Conformer 2. Also, the metal ions in Conformer 1 are somewhat less variable. In both molecules, there is flexibility around the 4WJ.

Compared to Conformer 1, Conformer 2 exhibits even more pronounced changes. While both metal ions remain bound, L3 is shifted away from P1.1. As a consequence, the M_A,Mg_ site in Conformer 2 has lost one metal contact, that to U52 in L3, consistent with this loop’s shift. The M_B,Mn_ site is concomitantly moved by 2 Å relative to its position in Llac-Mn and Conformer 1 (**Fig. 1d**). Three of its metal contacts are lost, specifically the inner-sphere contacts to the phosphates of C51, U52, and G9, which are now too far to interact at 6.8, 5.8, and 4.4 Å, respectively (**Fig. 2b**). Two of these lost contacts, U52 and G9, are from phosphates that bridge the two metal sites. Interestingly, the electron density is sufficiently clear to suggest that A48 in the binding site is flipped relative to its previous orientation, now orienting its N1, rather than N7, toward M_B,Mn_. Consistent with this generally disorganized architecture, the temperature B-factors for L3 are higher in Conformer 2 than Conformer 1 (**Fig. 2c,d**), indicating increased flexibility. Strikingly, this includes larger flexibility around P1.1, suggesting a possible link between the flexibilities of these two key regions. Taken together, our crystal structures provide first evidence that lost contacts between the metal ions, L1, and a partially unfolded L3 can translate into disorder in the distal P1.1 helix, which provides a basis for riboswitching.

### MD simulations reveal effects of M_A,Mg_ and M_B,Mn_ inner-sphere contacts on L3 stacking

To provide deeper insights into the RNA dynamics associated with metal identity, we performed 29 atomistic molecular dynamics (MD) simulations, equivalent to a total of 53 μs of real-time (see **Table 1**, **Methods**), with different metal ions in the M_A,Mg_ and M_B,Mn_ sites and starting either from Xory-Mn Conformers 1 or 2 (as reported here) or from Llac-Mn. As expected, given their necessarily limited timescales compared to experiments^22,23^, we found that our simulations generally maintained quite stable inner-sphere contacts for divalent metal ions, including their coordination with water molecules (**Fig. 3a-c**). To accelerate these dynamics we performed additional MD simulations that replaced one or both divalent ions at the M_A,Mg_ and M_B,Mn_ ion binding sites with monovalent K^+^ ions (**Table 1**). Importantly, while the divalent-to-monovalent replacement affects both the thermodynamics and kinetics of the inner-sphere contacts of the metal ions, their relative thermodynamic stability allows monovalent first-shell ligands to model the functional behavior of divalents on a shorter timescale^22,23^. Accordingly, even these more dynamic simulations maintained the global architecture of the riboswitch except for local perturbations of L3 and a variable P2-P4 helical angle (**Supplementary Fig. 4** and **Supporting Information**).

**Figure 3.**
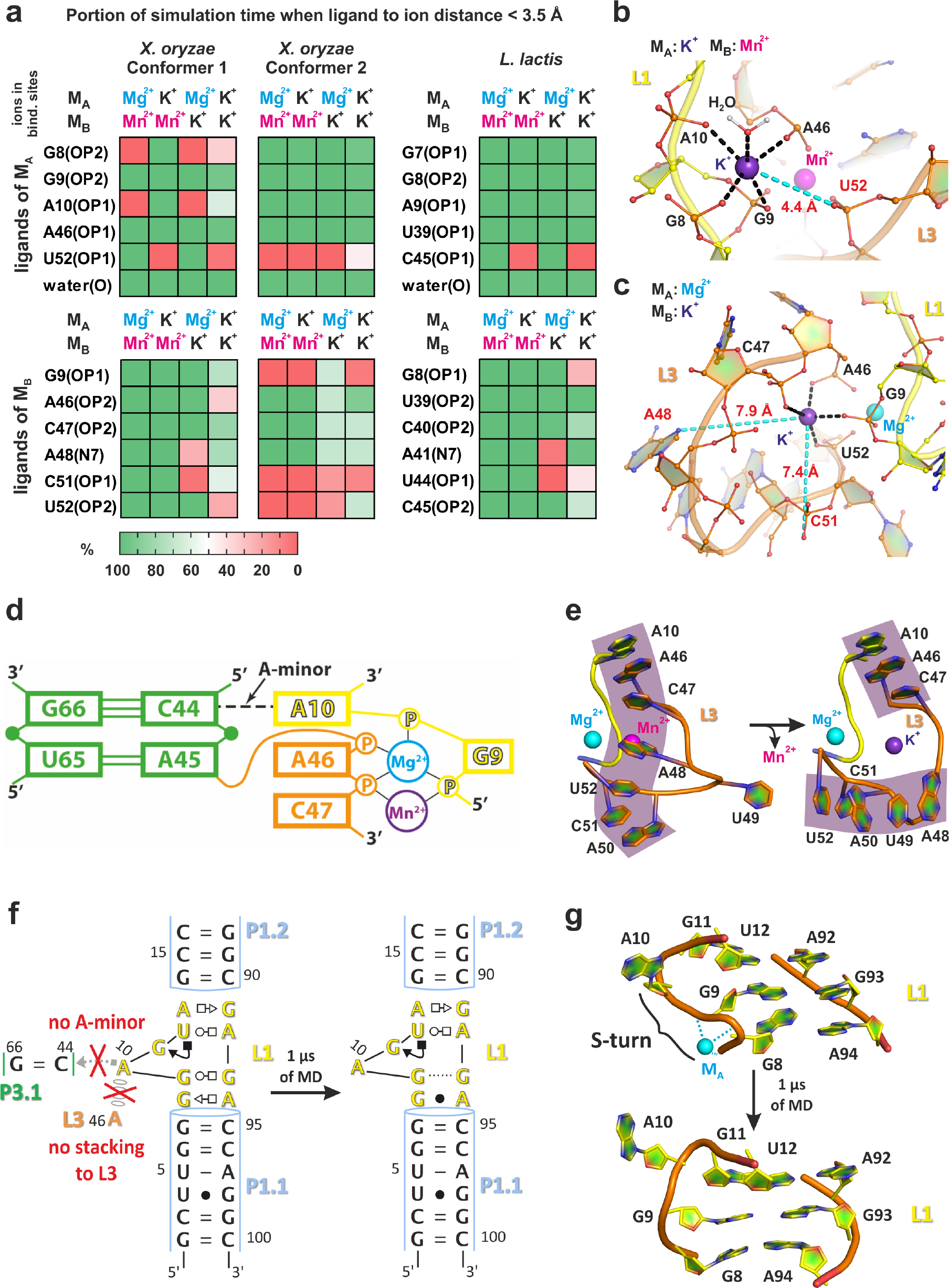
Structural dynamics of inner-shell ligands of the M_A,Mg_ and M_B,Mn_ sites, L3 and L1 loops as revealed by MD simulations. (**a**) The population of native inner-shell contacts in the M_A,Mg_ and M_B,Mn_ sites in percent as a function of the type of ion in these sites. A representative structure of the (**b**) M_A,Mg_ and (**c**) M_B,Mn_ site as typically observed in MD simulations with a K^+^ ion occupying the site. The most labile inner-shell contacts are depicted in cyan. (**d**) 2D representation of the most stable part of the L1-L3 tertiary interaction that is likely anchored by the A-minor interaction and related A10|A46 stacking. (**e**) Partial unfolding of the stacking pattern in the L3 loop observed upon replacement of Mn^2+^ with K^+^ in the M_B,Mn_ site. (**f**) The loss of native G9-G93 *trans* Watson-Crick/Hoogsteen and G8-A94 *trans* Sugar-Edge/Hoogsteen base pairing, and (**g**) loss of the S-turn backbone conformation involving a phosphate notch forming part of the M_A,Mg_ ion binding site in response to loss of the L1-L3 tertiary interaction with its A10…G66=C44 A-minor interaction.

**Figure 4.**
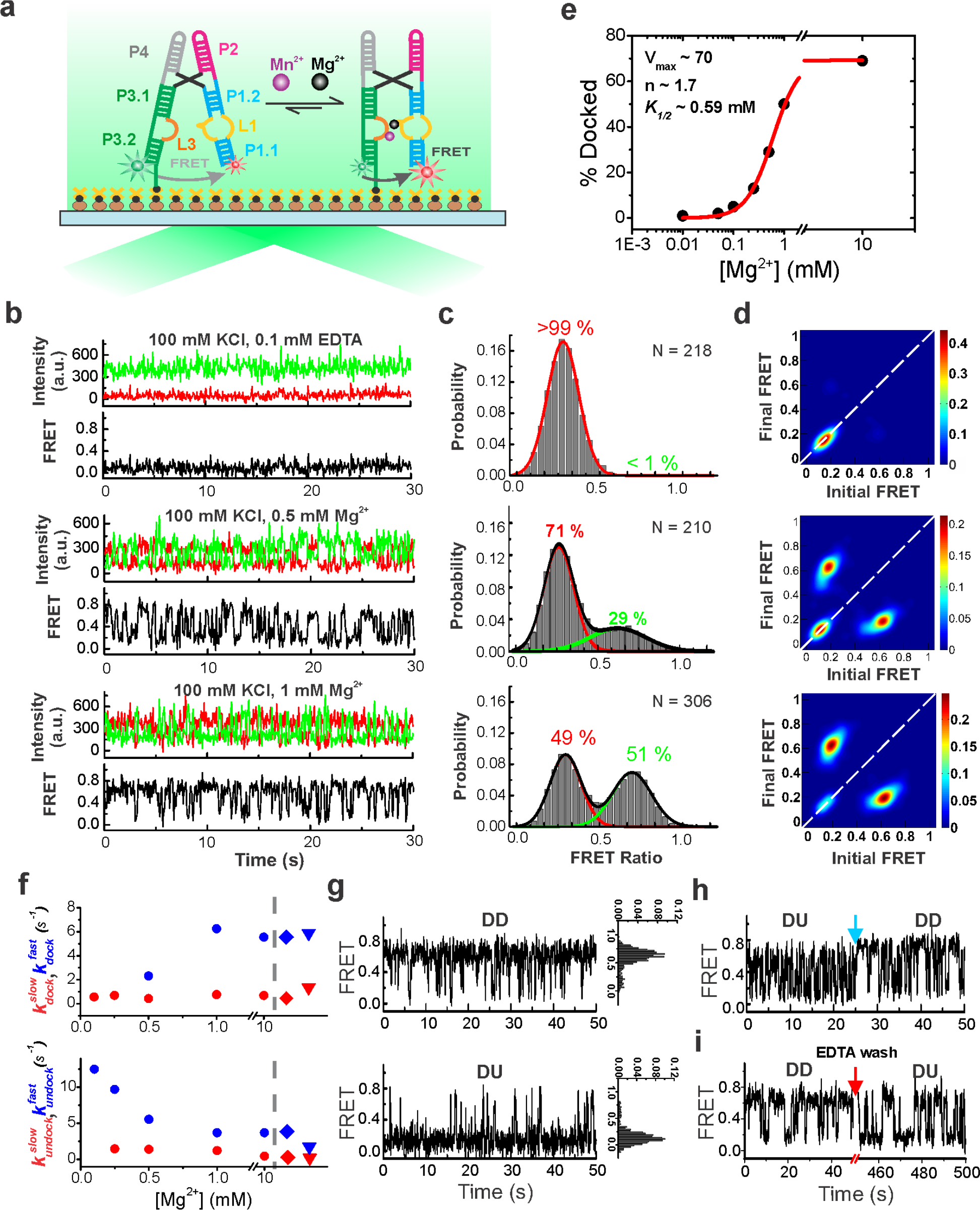
smFRET analysis of the WT *Xory* Mn^2+^ riboswitch. (**a**) Schematic of the smFRET experiment using TIRFM indicating the fluorophore labeling positions on the riboswitch. (**b**) Representative smFRET traces under different buffer conditions (top – bottom): absence of divalents (+0.1 mM EDTA), 0.5 mM Mg^2+^ and 1 mM Mg^2+^, respectively. Green, Cy3; Red, Cy5; Black, FRET. (**c**) Population FRET histograms showing the equilibrium distribution of two FRET states under the conditions of panel (**b**). Gaussian peaks for the low- and high-FRET states are shown in red and green, respectively with the cumulative fit shown in black. Reported are the percentages of FRET states at equilibrium, as well as the number of molecules N analyzed. (**d**) TODPs showing the ‘static’ and ‘dynamic’ trances as ‘on-diagonal’ and ‘off-diagonal’ heat map contours, respectively. The color code indicates the fraction of each population. (**e**) Fraction of the high-FRET state as a function of Mg^2+^ concentration, fit with a standard Hill equation (black). (**f**) Kinetics of structural dynamics as a function of Mg^2+^ and Mn^2+^ concentration. The diamond symbols represent rates in 1 mM Mg^2+^ and 0.1 mM Mn^2+^ while the triangle symbols represent rates in 0.1 mM Mn^2+^ alone. (**g**) Exemplary dynamic docked (DD, top) and dynamic undocked (DU, bottom) traces in the presence of 1 mM Mg^2+^. FRET histograms for the individual traces are shown on the right. (**h**) Rare examples of interconversion between different kinetic regimes showing dynamic heterogeneity. The blue arrow indicates the time of switching between the two kinetic regimes. (**i**) Exemplary trace showing conversion of a DU trace to DD trace upon chelation of Mg^2+^ with EDTA (indicated by the red arrow) and reintroducing 1 mM Mg^2+^.

Upon closer inspection, we found that the inner coordination sphere of monovalent ions in both the M_A,Mg_ and M_B,Mn_ sites were significantly more stable in simulations where the second binding site was occupied by a divalent rather than monovalent ion (**Fig. 3a**). This observation suggests that binding of the divalent ion into one binding site may help pre-organize and stabilize the second binding site, consistent with the notion of cooperativity between M_A,Mg_ and M_B,Mn_. Furthermore, while some inner-sphere contacts remained very stable, other contacts were lost upon replacement with a monovalent (**Fig. 3a-c**). Notably, the bridging inner-sphere contacts of the U52 phosphate with the M_A,Mg_ ion and those of the C51 and G9 phosphates with the M_B,Mn_ metal were found to be most labile across all simulations (**Fig. 3a**, **Supplementary Table 2** and **Supplementary Figs. 4**,**5**). These are the same inner-sphere contacts found to be lost in the crystallized Conformers 1 and 2 compared to the Llac-Mn, supporting the notion that the Conformers are relevant *en route* to (un)folding of the metal ion binding sites. In contrast, the inner-shell contacts of the A46 phosphate with the ions in both M_A,Mg_ and M_B,Mn_, as well as those of the G9 phosphate with the ion in M_A,Mg_ and of the C47 phosphate with the ion in M_B,Mn_, were stable across all simulations (**Fig. 3a**, **Supplementary Table 2** and **Supplementary Figs. 4,5**). These latter interactions are likely stabilized by coupling of the A-minor interaction of A10 to the G66-C44 base pair and associated stacking of A10 on the neighboring A46. All these interactions are parts of the L1-L3 tertiary docking contact and remained stable across all simulations (**Fig. 3d**).

**Figure 5.**
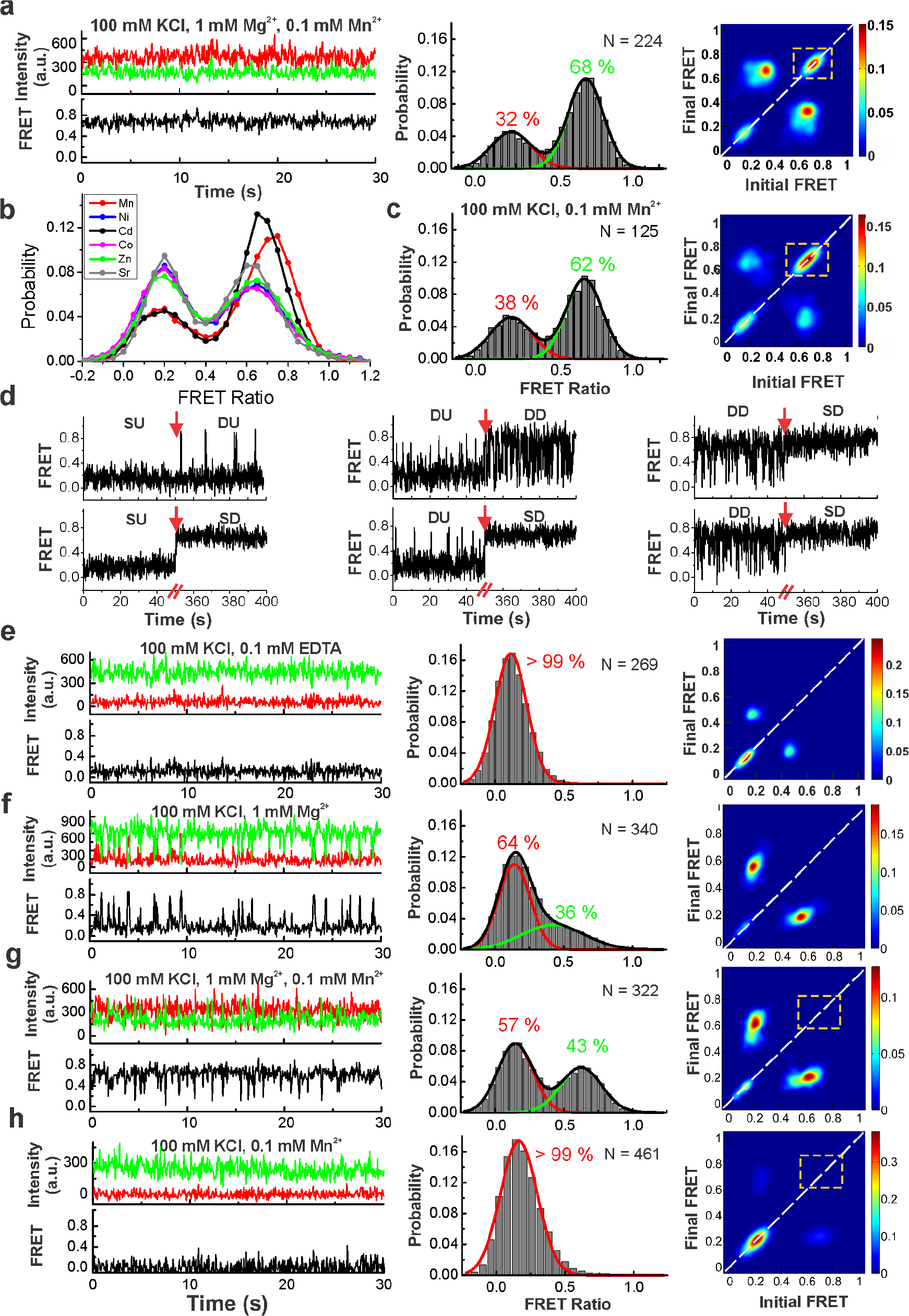
Effect of Mn^2+^ and A48U mutation on the folding of the riboswitch. (**a**) Representative smFRET trace, FRET histogram and TODP for the WT riboswitch in the presence of 0.1 mM Mn^2+^ and 1 mM Mg^2+^. The stable ‘static’ docked (SD) conformation appears only in the presence of Mn^2+^, as highlighted by the dashed yellow box in the TODP. (**b**) FRET histograms for the WT riboswitch at 0.1 mM concentration of different transition metals in the presence of 1 mM Mg^2+^. Over 150 molecules were analyzed for each histogram. The corresponding TODPs are shown in **Supplementary Fig. 16**. (**c**) FRET histogram and TODP with SD population highlighted by dashed yellow box for the WT riboswitch in the presence of 0.1 mM Mn^2+^ alone. (**d**) Example smFRET traces for the WT riboswitch showing transitions between SU, DU, DD and SD traces upon addition of 0.1 mM Mn^2+^. Representative smFRET traces, FRET histograms and TODPs for the A48U mutant riboswitch under: (**e**) no divalents (+0.1 mM EDTA), (**f**) 1 mM Mg^2+^ alone, (**g**) 1 mM Mg^2+^ and 0.1 mM Mn^2+^, and (**h**) 0.1 mM Mn^2+^ alone. The dashed yellow box highlights the absence of SD conformations for the A48U riboswitch in the presence of Mn^2+^, in contrast to the WT riboswitch.

We also found the dynamics of the entire L3 loop to be sensitive to the type of ion in the M_B,Mn_ site. When Mn^2+^ in the M_B,Mn_ site was replaced with K^+^, the flexibility of the L3 loop was increased so that the loop populated various conformations with broken stacking patterns (**Fig. 3b** and **Supporting Information**). These perturbations originated from the weakened inner-sphere contact of the A48(N7) nitrogen to the M_B,Mn_ metal, suggesting that the L3 loop is involved in direct sensing of the Mn^2+^ ion by forming a linchpin to support a tight stacking patterns only in the presence of the native Mn^2+^ (**Fig. 3e**).

### Stability of the SRL-like conformation of L1 loop requires an A-minor interaction

To complement our probing of structural dynamics in the docked state with those in the undocked state, we performed additional MD simulations of the segment consisting only of P1.1, P1.2 and L1. We started these “undocked” simulations from either of the two Xory-Mn Conformers and Llac-Mn (**Table 1**), with the aim to reveal their structural dynamics in the absence of the L1-L3 docking contact. While the L1 loop in the context of the entire riboswitch with its docked L1-L3 interaction always populated the SRL-like, two of our three “undocked” simulations lost this motif (**Figs. 3f,g**, **Supplementary Fig. 8**, and **Supporting Information**). In particular, in one of the simulations we observed loss of the G9-G93 *trans* Watson-Crick/Hoogsteen and G8-A94 *trans* Sugar-Edge/Hoogsteen base pairing^21^ as well as the S-turn^24^ that forms part of the M_A,Mg_ site (**Fig. 3f,g**). The former two base pairs are coaxially stacked on the P1.1 stem, suggesting that their loss may highlight the beginning of a transduction path by which P1.1 could become destabilized.

### smFRET reveals an undocked conformation that transiently docks upon Mg^2+^ addition

To probe the global structural dynamics in the presence of Mg^2+^ and Mn^2+^, we used smFRET to monitor fluorophores positioned on the distal legs of the Xory riboswitch (**Fig. 4a**, **Supplementary Fig. 13**, and **Methods**). smFRET traces at 100 mM KCl in the absence of any divalent metal ions showed a stable low-FRET value of ~0.1 without global dynamics (**Fig. 4b**), with a population FRET histogram displaying a single peak centered on ~0.13±0.10 (mean ± standard deviation) (**Fig. 4c**). The non-dynamic nature of the traces is also evident as an ‘on-diagonal’ contour centered at ~0.13 in the transition occupancy density plot (TODP), which shows initial vs. final FRET value transitions for the population of molecules as a heat map (**Fig. 4d**). This FRET value corresponds to an estimated distance of ~74 Å between the two fluorophores and suggests an extended, stably undocked (SU) conformation where the two RNA legs are distal and do not interact, unlike the docked crystal structure (**Fig. 1b**).

Addition of Mg^2+^ up to 0.1 mM did not result in significant changes in the FRET histograms since almost all traces remained in the SU conformation, with ~3 % of them showing brief excursions into a higher ~0.6-FRET state (**Supplementary Fig. 14**). Further raising the Mg^2+^ concentration resulted in more dynamic traces transiently adopting this high-FRET state, accompanied by a corresponding decrease in the population of SU traces (**Fig. 4b-d**, **Supplementary Fig. 14**). At a near-physiological concentration of 1 mM Mg^2+^, the time- and population-averaged distribution between low- and high-FRET, with mean FRET values of ~0.15±0.11 (49 %) and 0.63±0.14 (51 %), respectively, became almost equal (**Fig. 4c**). A FRET value of 0.63 corresponds to a distance of ~49 Å between the two labeled RNA arms, similar to the distance observed in the crystal structures, suggesting adoption of the compact ‘docked’ conformation. Reaching 10 mM Mg^2+^, the fraction of this docked conformation further increased and saturated at ~69 %, with a sigmoidal Mg^2+^ concentration dependence that fit well with a Hill equation to yield a half-saturation point of *K*_*1/2*_ ~ 0.6 mM and a cooperatively coefficient of n = 1.7 (**Fig. 4e**). These data demonstrate that the Mn^2+^ riboswitch adopts an extended SU conformation in the absence of divalents, which increasingly samples transient docked conformations upon a rise in Mg^2+^ concentration.

At a low Mg^2+^ concentration of 0.1 mM, single-exponential kinetics were observed with a slow docking rate constant, *k*_*dock*_ ~0.56 s^−1^, and a fast undocking rate constant, *k*_*undock*_ ~12.5 s^−1^ (**Fig. 4f** and **Supplementary Fig. 15**). Further increasing the Mg^2+^ concentration to 1 mM resulted in the emergence of double-exponential kinetics in both *k*_*dock*_ and *k*_*undock*_. The docking kinetics exhibited 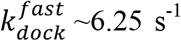 and 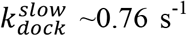, while the undocking kinetics displayed 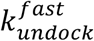 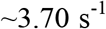 and 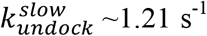 (**Fig. 4f** **and** **Supplementary Fig. 15**). The TODP further shows that a majority (82 %) of traces are dynamically transitioning between the two FRET states, as highlighted by dominant ‘off-diagonal’ contours, while only a small fraction (~18 %) remains in the stable low-FRET state (**Fig. 4d**). The double-exponential kinetics arises from two distinct populations: ‘dynamic docked’ (DD) and ‘dynamic undocked’ (DU) traces corresponding to molecules residing largely in the docked and undocked states, respectively. (**Fig. 4g**). Among the dynamic traces, ~65 % were DD while ~35 % were DU traces. As observed for other RNAs that undergo docking of two adjacent helical arms^25–31^, the heterogeneity observed in the population was largely static and molecular behaviors interconverted only rarely (< 2% of traces) over the experimental timescale (5-10 min) (**Fig. 4h**). Interconversion between the DU and DD behaviors was observed more readily, however, when first chelating, then reintroducing Mg^2+^ (**Fig. 4i**), suggesting that they represent kinetically trapped conformations on a deeply rugged folding free energy landscape^32,33^.

### Sub-millimolar Mn^2+^ uniquely yields a stably docked riboswitch

We next asked what specific effect Mn^2+^ has on the folding of the riboswitch. In the presence of 1 mM Mg^2+^, addition of a low concentration of 0.1 mM Mn^2+^ resulted in the appearance of a unique population of stably docked (SD, ~43%) traces residing in the high-FRET state for >30 s (*k*_*undock*_ <0.03 s^−1^) before photobleaching (**Fig. 5a**). Accordingly, the FRET histogram showed two peaks with mean FRET values of 0.17±0.14 and 0.69±0.12 and an increased ~68 % population of the docked conformation (**Fig. 5a**). The SD population is evident in the TODP as a new ‘on-diagonal’ contour centered on the ~0.7-FRET value (**Fig. 5a**). Dynamic traces under these conditions again showed double-exponential behavior and were similar to the kinetics at 1 mM Mg^2+^ alone, with 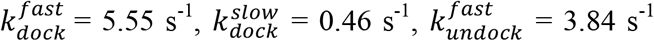 and a ~5-fold slower 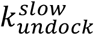 of 0.23 s^−1^. Notably, in the presence of Mn^2+^, most of the dynamic traces now showed DD character. These data demonstrate that Mn^2+^ binding generally stabilizes the docked conformation while uniquely enabling a new, stably docked SD state. FRET histograms further showed that out of a variety of divalent metal ions tested, only Cd^2+^ is effective in promoting docked conformations over just 1 mM Mg^2+^ alone; Ni^2+^, Co^2+^, Sr^2+^ or Zn^2+^ had little effect (**Fig. 5b**, **Supplementary Fig. 16**). Interestingly, in the absence of Mg^2+^, while 0.1 mM Mn^2+^ alone led to the appearance of DD and SD traces with ~62 % docked population (mean FRET 0.67±0.12) (**Fig. 5c**), 0.1 mM of Ni^2+^, Co^2+^, Sr^2+^ or Zn^2+^ did not affect SU traces and Cd^2+^ had only a small effect in promoting DD traces. These results suggest that while the *Xory* riboswitch has some degree of plasticity in recognizing ligands, in a background of Mg^2+^ it preferentially recognizes Mn^2+^ and – to a lesser extent – Cd^2+^.

To probe how the kinetically distinct SU, DU and DD traces respond to Mn^2+^, we observed the same set of molecules at 1 mM Mg^2+^ before and after the addition of Mn^2+^. We found that all three populations respond to Mn^2+^ and are capable of forming the stably docked (SD) population (**Fig. 5d**). In particular, the SU traces converted into DU, DD, and SD conformations with similar probabilities, suggesting that they are correctly folded with metal sensing sites poised to bind ligand. By comparison, a majority of DD traces converted into SD traces whereas DU traces adopted DD and SD behavior upon Mn^2+^ addition. Only a small fraction of traces showed no response at low Mn^2+^ suggesting that they may be misfolded.

### Mutation of the conserved adenosine A48 results in complete loss of the SD conformation

The highly conserved discriminator base A48 in L3 is positioned to confer Mn^2+^ specificity via its N7 and also helps maintain L3 in a stacked conformation. We therefore tested the effect of a single A48U mutation on folding and Mn^2+^ sensing of the riboswitch. Similar to the WT, at 100 mM KCl without divalents, smFRET traces of the mutant riboswitch showed the SU population with a mean FRET value of 0.11±0.12 (**Fig. 5e**). At 1 mM Mg^2+^, we observed dynamic traces with excursions into the docked higher FRET states, and the FRET histogram showed a major (64 %) 0.14±0.11 low-FRET peak and a minor (36 %) broad 0.52±0.21 mid-FRET peak that corresponds to more extended docked conformations (**Fig. 5f**). Of note, 100% of the dynamic mutant traces showed DU character with only brief excursions into the docked conformations (**Fig. 5f**). As a result, we observed fast single-exponential undocking with *k*_*undock*_ ~6.67 s^−1^ while *k*_*dock*_ was double-exponential with a major 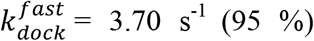 and a minor 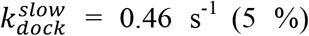 (**Supplementary Fig. 17**). The TODPs show SU behavior in the absence of divalents, whereas both SU and DU behaviors are observed at 1 mM Mg^2+^ (**Fig. 5e,f**).

Next, we asked whether the mutant riboswitch can still respond to Mn^2+^. In the presence of 1 mM Mg^2+^ and 0.1 mM Mn^2+^, most smFRET traces were dynamic and showed DD behavior, while only a small fraction remained in the SU state, similar to WT (**Fig. 5g**). FRET histograms show a similar ~0.14±0.13 (57 %) low-FRET undocked state but display a docked state now with a higher mean-FRET value of 0.63±0.15 and a larger ~43 % population (**Fig. 5g**), compared to the 1 mM Mg^2+^ only condition. . The kinetics under these conditions were double-exponential with a 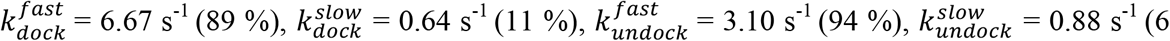 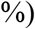 (**Supplementary Fig. 17**). Importantly, the mutation caused a striking loss of SD traces, as evident from the complete absence of the on-diagonal ~0.7 FRET contour in the TODP (**Fig. 5g**), in stark contrast to the WT. In addition, the mutant interestingly loses the WT’s ability to sample the docked conformations in the presence of 0.1 mM Mn^2+^ alone (**Fig. 5g**); almost all smFRET traces at 0.1 mM Mn^2+^ are in the SU conformation, with the histogram showing a major single peak around 0.19±0.13 FRET, further corroborated by a single on-diagonal ~0.2 FRET contour in the TODP (**Fig. 5h**). These data demonstrate that the conserved discriminator A48 is essential for Mn^2+^ inducing a stably docked riboswitch conformation. The presence of Mg^2+^ partially rescues the loss of Mn^2+^ sensing by the mutant, yet does not restore its ability to form the SD conformation. Possibly this reflects the ability of Mg^2+^ to bind at M_B_ in A48U, though with weaker affinity, via the uridine O4, as suggested in the A41U *L. lactis* structure^12^.

## DISCUSSION

Using a combination of X-ray crystallography, MD simulations and smFRET, our work here sheds light on how the binding of a single Mn^2+^ ion can modulate the stability of the distant switch helix through a conformational cascade that couples local with global structural dynamics. While previous structures revealed the local Mn^2+^ recognition mechanism of *yybP-ykoY* RNA^12^, the new structures reported here underscore the conformational plasticity around the metal binding core. Importantly, the structural differences observed between the two conformers in the crystal lead us to posit that Conformer 2 with L3 partially unfolded around M_B,Mn_ is likely an intermediate between the fully Mn^2+^-bound state and the empty M_B,Mn_ state observed in the *E. coli* Mn^2+^-free structures^12^. We hypothesize that the loss of phosphate contacts from C51, U52, and G9 to Mn^2+^ in Conformer 2 reflects their relative weakness as supported by MD simulations and likely represents a discrete step in Mn^2+^ binding/unbinding. Conversely, the variation in M_B,Mn_ site coordination we observe could represent inherent flexibility at this site, allowing it to recognize Mn^2+^ even if the metal ion is not fully dehydrated. This flexibility is also seen in the recent Cd^2+^-bound structures (see **Supporting Information**).

Our MD simulations show that the SRL-like motif in L1 is inherently unstable in the L1-L3 undocked state, but is stabilized by the tertiary A-minor interaction of A10 with P3.1. Similarly, L1-L3 docking stabilizes L3 in a stacked conformation by direct stacking of A10 on top of it. Thus, a stable conformation of L1 and therefore P1.1 is intimately linked to the stability of L3 and the tertiary A-minor interaction. We suggest that the SRL-like motif acts as a molecular switch that links the Mn^2+^-dependent increase in docking observed by smFRET to stabilization of P1.1.

Using smFRET we further identify a globally undocked conformation at physiological Mg^2+^ concentration that remains in dynamic equilibrium with the compact docked conformations. In fact, the observation of pairs of stably and dynamically docked and undocked states with double-exponential docking and undocking kinetics supports the existence of at least two undocked and docked dynamic (in addition to the static SU and SD) conformations, with similar FRET values and thus global folds, but likely different local conformations around the ligand sensing core. These conformations are possibly related to Conformers 1 and 2 in the crystal structure. We further find that SD conformations are formed only in the presence of sub-millimolar Mn^2+^ or the related soft transition metal Cd^2+^, and lost upon a single mutation of the invariant Mn^2+^-sensing adenine A48.

Our structural snapshots, MD simulations and smFRET data support the model shown in **Fig. 6**. In the absence of any divalent ions, the *Xory* riboswitch adopts an extended conformation in which the two legs are distal and rarely interact, as shown by smFRET. Under these conditions, the metal-binding sites are likely unstructured and their interaction only transiently formed by L1-L3 contacts, as suggested by rare smFRET transitions, MD simulations as well as previous ligand-free structures^12^. Our MD simulations in particular show that the A-minor interaction formed by A10 of L1 loop with the GC base pair of stem P3.1 can at least transiently form even in the absence of divalents. The formation of this A-minor interaction helps pre-organize the M_A,Mg_ binding site. Under physiological (1 mM) concentrations of Mg^2+^, this ion binds first at the M_A,Mg_ site, allowing the riboswitch to sample dynamic folded conformations where the two legs are brought together via metal-mediated interaction between loops L1 and L3. These dynamic folded states then allow for local interactions mediated by L3, forming a pocket of high negative charge potential poised to sense Mn^2+^. Mg^2+^ binding at M_A,Mg_ therefore facilitates binding of Mn^2+^ at M_B,Mn_ in cooperative fashion, as shown by smFRET at the global level and further visualized by atomistic MD simulations. Capture of Mn^2+^ at site M_B,Mn_ then acts as the final linchpin to hold the two legs together most stably. This compact docked conformation with a SRL-like fold of L1 and continuous coaxial stacking between P1.2, L1 and P1.1 stabilizes the remote P1.1 ‘switch’ helix and prevents the strand invasion required to form the terminator hairpin, instead promoting transcriptional read-through.

**Figure 6.**
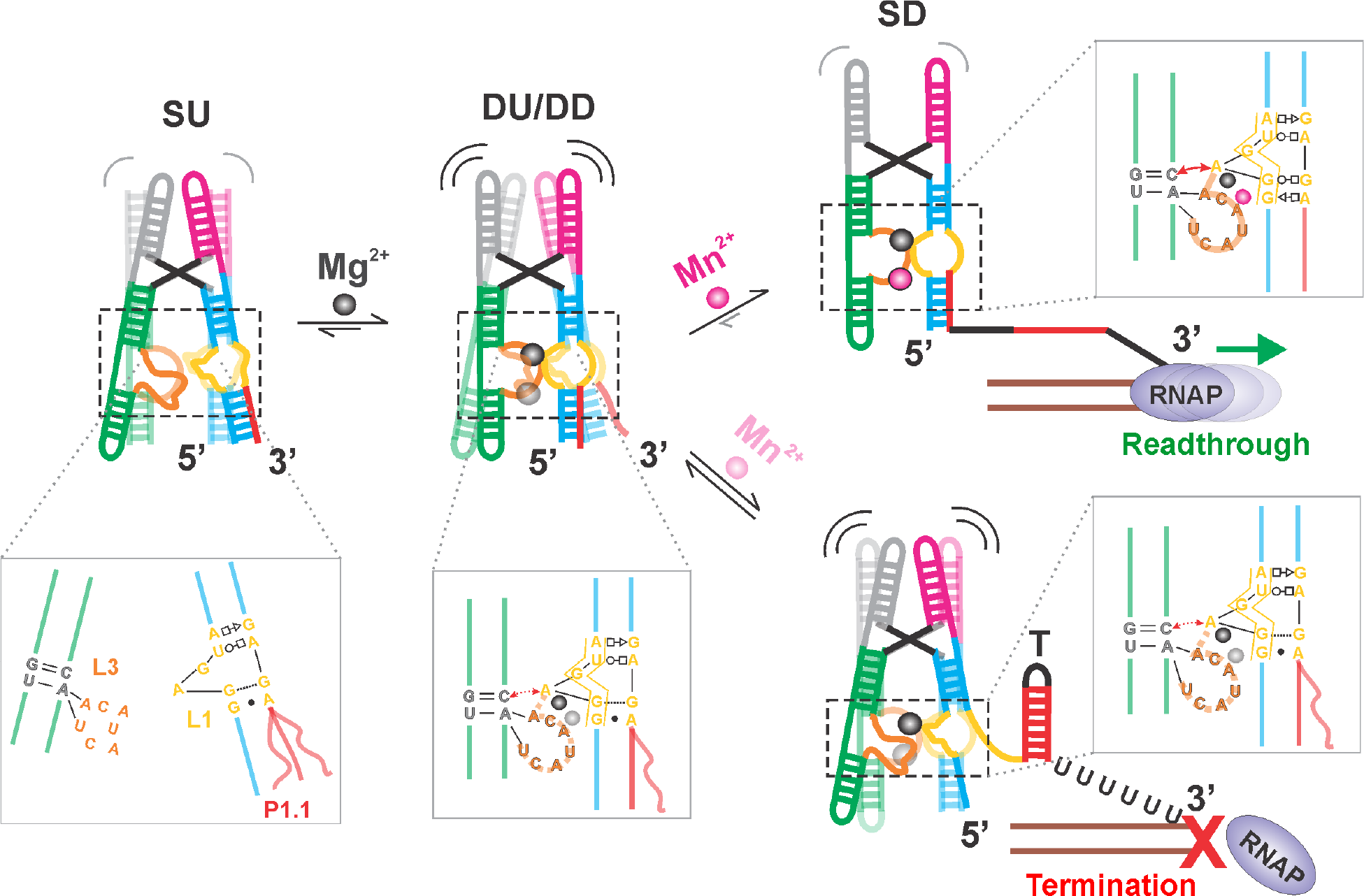
Local-to-global signal transduction pathway in the core of the *Xory* Mn^2+^ sensing riboswitch. The *Xory* riboswitch exists in an ‘X-shaped’ extended SU conformation in the absence of Mg^2+^, with very rare transitions into docked ‘H-shaped’ conformations. In the SU state, the legs are far apart in the absence of a stable A-minor tertiary interaction and loops L3 and L1 are less structured under these conditions. In the presence of physiological Mg^2+^ concentrations (mM), the riboswitch samples docked conformations with distinct kinetics, indicative of differences in L3 stacking as observed in our crystal structures. L1 adopts an S-turn conformation (as indicated by the yellow box in the inset) and L3 is partially stacked under these conditions, aided by the presence of Mg^2+^ ions in M_A,Mg_ and transiently in M_B,Mn_. In the presence of sufficient Mn^2+^, binding of the metal ion by A48 reinforces L3 stacking and stability of the A-minor interaction. This maintains L1 in a rigid non-canonical SRL-like conformation that enables co-axial stacking between P1.2 and the switch helix P1.1. In turn, this cascade of interactions stabilizes P1.1, thereby preventing strand invasion to favor formation of the antiterminator required for transcriptional read-through.

Contrasting with the Mn^2+^ riboswitch, the NiCo riboswitches, which cooperatively sense the transition metals Ni^2+^ or Co^2+^, resemble the overall ‘H’ shaped architecture of the Mn^2+^ riboswitch but with a tight 4WJ that lacks an analogous tertiary docking interface. Accordingly, the NiCo riboswitch appears to acts through a distinct mechanism utilizing four bound metal ions to weave together a network of interactions between the interhelical residues that stabilize the 4WJ and so prevent formation of a terminator via strand invasion^34^. Similarly, the Mg^2+^ sensing M-box riboswitch adopts an architecture with three parallel co-axially stacked helices that are brought together by the binding of six Mg^2+^ ions, leading to formation of a terminator^35^. Therefore, both the NiCo and Mg^2+^ sensing riboswitches achieve gene regulation by employing multiple metal ions that directly interact with and stabilize the ‘switch’ helix. The Mn^2+^ riboswitch is thus unique in requiring only a single metal ion that does not make any direct contacts with switch helix P1.1. By combining high-resolution structural information with insights into both atomistic and global dynamics, our study outlines the mechanism by which a ligand as small as a single divalent metal ion couples local with global structure to give Mn^2+^ the ability to influence RNA folding and fine-tune gene expression by cooperating with a high background of Mg^2+^. The general lessons revealed here of how a ligand-binding signal can be transduced across an RNA are likely to become a recurring theme among riboswitches where the ligand represents a distal structural linchpin for the ‘switch’ helix.

## Acknowledgements

This work was supported by NIH R01 grants GM062357 and GM118524 to N.G.W., by project SYMBIT reg. number CZ.02.1.01/0.0/0.0/15_003/0000477 funded by the ERDF (M.J. and J.S.), by grant 18-25349S (P.B., P.K.) from the Grant Agency of the Czech Republic, and a student project of Palacky University Olomouc IGA_PrF_2018_032 (M.J.). We thank Qian Hou for help with some experiments. This work included research conducted at NE-CAT beamlines, which are funded by the National Institute of General Medical Sciences from the National Institutes of Health (P30 GM124165). The Pilatus 6M detector on 24-ID-C beam line is funded by a NIH-ORIP HEI grant (S10 RR029205). This research used resources of the Advanced Photon Source, a U.S. Department of Energy (DOE) Office of Science User Facility operated for the DOE Office of Science by Argonne National Laboratory under Contract No. DE-AC02-06CH11357.

## ONLINE METHODS

### RNA preparation and crystallization

RNA was cloned, transcribed, purified, refolded with 2.5 mM Mn^2+^, and screened for crystallization as previously described^12^. The *X. oryzae* aptamer domain sequence was modified to improve the chances of crystallization, including replacing the terminal loops in variable regions with GAAA tetraloops and adding a GG at the beginning of the sequence to increase T7 RNA polymerase efficiency^36^. We also removed a single unconserved U flip-out near the base of P1.1. In this construct, a native CA dinucleotide at the four-way junction was omitted from the sequence (**Fig. 1a**), possibly aiding crystallization. Initial crystal hits were obtained using the Nucleic Acid Mini-Screen (Hampton Research). This RNA crystallized at 0.2 mM in 10% MPD, 40 mM Na cacodylate (pH 7), 12 mM spermine tetrahydrochloride, 80 mM SrCl_2_, and 20 mM MgCl_2_ after 2-3 months at 18 °C.

### X-ray data collection and structure building

A 2.85 Å resolution X-ray diffraction dataset was collected at NE-CAT beamline 24-ID-C at the Advanced Photon Source (Argonne, IL), at 0.769 Å wavelength in order to use the Strontium K-edge for phasing. An additional dataset was collected at a remote wavelength (0.7749 Å) for SIRAS. Datasets were processed by XDS^37^ as part of NE-CAT’s RAPD pipeline. An initial solution was achieved by molecular replacement using the central region of the *L. lactis* structure^12^ followed by finding portions of external helices^38^. This partial solution was then used for MR-SAD in AutoSol in the Phenix suite^39,40^ to find Sr sites and obtain an initial electron density map. Since this dataset was collected near the Sr K-edge wavelength (0.769 Å), we could observe anomalous signal from Sr^2+^ ions in the crystal. At 4 σ (a relatively low threshold), there was modest signal at the Conformer 2 M_A,Mg_ site, suggesting only partial occupancy by Sr^2+^ (**Fig. 2b**). However, there is no occupancy seen at any of the other M_A,Mg_ or M_B,Mn_ sites, despite there being external sites in the lattice with stronger anomalous signal (**Supplementary Fig. 1**). This suggests that the Conformer 2 M_B,Mn_ site, despite lacking several contacts, can still distinguish between metals. It possibly discriminates by geometry; the Sr^2+^ ionic radius is 2.4 Å, compared to the smaller Mn^2+^ (2.2 Å) or Mg^2+^ (2.1 Å). Mg^2+^ cannot be differentiated from Mn^2+^ at the wavelength and resolution of this structure. Alternating rounds of building in Coot^41^ and refinement with phenix.refine^42^ and intermittent re-phasing and density modification in AutoSol were used to build the final model (**Supplementary Table 1**).

### Molecular Dynamics Simulations

The crystal structures of Mn^2+^ sensing riboswitch from *X. oryzae* from this study (both Conformers 1 and 2) and from *L. lactis* (PDB ID 4Y1I^12^) were used as starting structures for MD simulations. We aimed to study the effect of divalents in the metal ion binding sites on the structural dynamics of the riboswitch, so we prepared a set of starting structures containing different ions (Mn^2+^, Mg^2+^, and K^+^) in the M_A,Mg_ and M_B,Mn_ sites (**Table 1**). In addition, we probed both for syn- and anti-conformations of A48 in Conformer 2 from *Xory* to verify refinement of the corresponding electron density as a *syn*-oriented nucleotide. Starting structures of the SRL-like motif lacking tertiary interactions with the rest of the aptamer were prepared from the crystal structures but entailing only the P1.1, P1.2 and L1 segments without divalents.

All MD simulations were carried out using pmemd.cuda (GPU code of AMBER 14 program package)^43,44^ with the ff99bsc0χ_OL3_ force field^45–47^. The simulation protocol was as follows. The structure with the RNA molecule and any ions in the M_A,Mg_ and M_B,Mn_ sites was immersed into a solvation box of SPC/E explicit water molecules^48^. We added KCl salt excess corresponding to 150 mM KCl. We used following ionic parameters: K^+^ (r = 1.593 Å, ε = 0.4297 kcal/mol^49^), Mg^2+^ (r = 1.5545 Å, ε = 0.00295 kcal/mol^50^), Mn^2+^ (r = 1.4060 Å, ε = 0.0167 kcal/mol^51^), and Cl^−^ (r = 2.711 Å, ε = 0.0127 kcal/mol^49^). Where indicated, we replaced one or both divalent ions in the M_A,Mg_ and M_B,Mn_ sites by K^+^ monovalents so that we probed for each specific starting structure four different ionic configurations in these ion binding sites (**Supplementary Table 2**). The solvated systems were equilibrated as follows. Geometrical optimization of hydrogen atoms was followed by optimization of waters and ions, while the position of the RNA molecule remained restrained. Subsequently, all RNA atoms were restrained and the solvent molecules with counter-ions were allowed to move during a 500-ps long MD run under NpT conditions (p = 1 atm, T = 298.16 K) to relax the total density of solvent surrounding the RNA. Next, the RNA molecule was relaxed by several minimization runs, with decreasing force constant of position restraints applied to the sugar-phosphate backbone. After full relaxation, the system was heated in two steps: The first step involved heating under NVT conditions for 100 ps, whereas the second step involved final density equilibration under NpT conditions for an additional 100 ps. The particle-mesh Ewald (PME) method for treating electrostatic interactions was used, and simulation were performed under periodic boundary conditions in the [NVT] ensemble at 298.16 K using a weak coupling Berendsen thermostat with a coupling time of 0.2 ps. The SHAKE algorithm, with a tolerance of 10^−5^ Å, was used to fix the positions of all hydrogen atoms, and a 10.0 Å cut-off was applied to non-bonding interactions. The hydrogen mass repartitioning was used to change the mass of hydrogen and neighboring atoms to allow a 4 fs integration step. In the production runs of the isolated SRL-like motif (P1.2, L1, and P1.1 segments only) we applied upper-wall positional restraints to the hydrogen bonds of the terminal base pair of the P1.2 stem (i.e., the G16=C89 base pair proximal to the P2 stem in the complete structure) to mimic the stabilization effect of coaxial stacking of this base pair on the P2 stem. A parabolic restraint potential was applied at heavy atom distances of the particular hydrogen bonds above 3.5 Å with a force constant of 2.5 kcal/mol/Å^2^. The simulation time for each studied system was at least 2 μs (**Supplementary Table 2**).

### Data analysis

All trajectories were analyzed with the Ptraj module of the AMBER package^44^ and the simulations were visualized using VMD^52^. B-factors were calculated as average mass-weighted fluctuation of each residue over the entire trajectory aligned to the stem closest to the residue. This guarantees that the B-factors are not influenced by global motions and reflect local flexibilities only. The stacking interactions were detected using the G-vector of an eRMSD metric and were annotated using baRNAba^53^.

### Single-molecule FRET RNA preparation

The RNA for smFRET studies was annealed from two synthetic RNA oligonucleotides (**Supplementary Fig. 13**), ordered with the indicated modifications from IDT. The native CA dinucleotide omitted for crystallization was present in the smFRET RNA. Oligonucleotide 1 has a 5’-Cy5 and oligonucleotide 2 has 5’-Cy3 and 3’ biotin-TEG. In this two oligonucleotide design of the smFRET construct, the donor fluorophore Cy3 and biotin-TEG were placed on the 5’ and 3’ ends of stem P1.1while the acceptor fluorophore Cy5 is attached to the 5’ end of P3.2, positioning the fluorophores on either end of the distal arms that dock. For specifically probing the docking interaction, we extended the stem P1.1 to prevent end fraying. The biotin on the 3’-end of P1.1 was used for immobilization of the RNA onto a microscope quartz slide and subsequent prism-based total internal reflection fluorescence microscopy (TIRFM, **Fig. 4a**). Optimal refolding of the RNA was achieved with a mixture of 1 μM oligonucleotide 1 and 1.5 μM oligonucleotide 2 in 20 mM HEPES pH 7.0, 50 mM KCl. In PCR tubes, samples were heated to 70 °C for 2 min, then 2 mM MgCl_2_ was added, samples were allowed to cool to RT for 10 min, then put on ice until further use. To check assembly, ~1 pmol of each oligonucleotide (1-2 μL) was added to 10 μL running buffer supplemented with 10% glycerol and loaded onto a non-denaturing 8% polyacrylamide gel run at 4 °C. A buffer of 0.5x TBE supplemented with 2 mM Mg(OAc)_2_ and 50 mM KCl was used in the gel and as the running buffer.

### smFRET data acquisition and analysis

A small microfluidic channel was made by sandwiching a cleaned quartz slide and a glass coverslip using double-side tape. Beforehand, two holes were drilled into the quartz slide and afterwards tubing was attached to them to act as inlet and outlet ports that were used for flowing buffer and introducing the RNA. The two RNA oligonucleotides containing the Cy3, Cy5 and biotin modifications were mixed at a concentration of 2 μM each in 50 mM Hepes, pH 7.2, 100 mM KCl (1x buffer). The RNA was heated at 90 °C for 2 min, left at RT for 30 s to cool down, followed by addition of 2 mM MgCl_2_ and slow cooling down to RT over 15 min. This annealed RNA stock was used for carrying out smFRET experiments, for which a 15-25 pM RNA solution was made by diluting the 2 μM annealed stock using 1x buffer.

100 μL of diluted RNA solution was flowed onto the quartz slide coated with biotinylated-BSA and streptavidin for immobilization and incubated for 2 min. Any unbound RNA was washed off using 1x buffer. This step also washes off small (low nM) amounts of residual Mg^2+^ present in the stock. For experiments in the absence of divalents, 0.1 mM EDTA was included in the 1x buffer to chelate any contaminating divalent ions. Mg^2+^ titration experiments were performed on the same slide after washing off the EDTA using 1x buffer. We included an enzymatic oxygen scavenging system (OSS) containing 50 nM protocatechuate-3,4-dioxygenase and 5 mM protocatechuic acid in 1x buffer to prolong the longevity of the fluorophores. In addition, 2 mM Trolox (6-Hydroxy-2,5,7,8-tetramethylchromane-2-carboxylic acid) was included in the imaging buffer to suppress photoblinking of the dyes.

All smFRET movies were acquired at ~16 Hz, unless otherwise specified, using a prism-based total internal reflection fluorescence (TIRF) microscope with an intensified CCD (IPentamax, Princeton Instruments) or sCMOS camera (Hamamatsu ORCA-Flash4.0 V3), essentially as previously described^54,55^. Cy3 on the RNA was excited using a 532 nm laser and the emission from both Cy3 (donor) and Cy5 (acceptor) were simultaneously detected side-by-side on the camera. Towards the end of all movies, the Cy5 was directly excited using a 640 nm laser to check for the presence of the acceptor (Cy5) fluorophore. This helps in distinguishing the low-FRET (~0.1 FRET) states we observe from ~0 FRET states due to the absence or photobleaching of Cy5 in Cy3-labeled molecules. Raw movies were analyzed using IDL (Research Systems) to extract the time traces for all spots in Cy3 and Cy5 channels. Single molecule traces were then visualized using Matlab and only those with a minimum combined intensity (Cy3 + Cy5 intensity) of 300, showing single-step photobleaching of the dyes, a signal-to-noise ratio of >3, and longer than 6 s were selected for further analysis. Selected traces were then background-subtracted to correct for cross-talk and (minimal) bleed-through. We calculated the FRET ratio as *I*_*A*_/(*I*_*A*_+*I*_*D*_), where *I*_*A*_ and *I*_*D*_ are the background corrected intensities of acceptor (Cy5) and donor (Cy3), respectively. FRET histograms were made using the first 50 frames (3 s) of all traces (at least 200) in a given condition and fit with a sum of Gaussians using OriginPro 8.5. For kinetic analysis, traces were idealized with a two-state model corresponding to undocked (low-FRET) and docked (high-FRET) states using the segmental k-means algorithm in QuB software as previously described^56,57^. Cumulative dwell-time histograms were plotted from all extracted dwell times and fit with single- or double-exponential functions using OriginPro 8.5 to obtain the lifetimes in the undocked (τ_undock_) and docked (τ_dock_) states. Rate constants of docking and undocking were then calculated as *k*_*dock*_ = 1/τ_undock_ and *k*_*undock*_ = 1/τ_dock_. For the double-exponential fits, kinetics were calculated similarly using both the short and long dwell lifetimes to obtain the fast and slow rate constants, respectively.

## Supplementary Information

**Table S1.** Crystallographic data and refinement statistics for the *X. oryzae* aptamer structure. Figures in parentheses indicate statistics for the highest resolution shell.

|  | **<u>X. oryzae-Mn<sup>2+</sup></u>** |
| --- | --- |
| Beamline | APS 24 ID-C |
| Wavelength (Å) | 0.7690 |
| Resolution range (Å) | 62.64-2.85 (2.95-2.85) |
| Space group | P 21 21 21 |
| Unit cell | 81.30 85.17 92.46<br>90.00° 90.00° 90.00° |
| Total reflections | 119014 (11355) |
| Unique reflections | 15512 (1510) |
| Multiplicity | 7.7 (7.6) |
| Completeness (%) | 99.8 (99.1) |
| Mean I/sigma(I) | 12.9 (0.66) |
| Wilson B-factor | 83.22 |
| R-merge | 0.118 (2.93) |
| R-meas | 0.127 (3.15) |
| CC1/2 | 1.00 (0.306) |
| CC* | 1 (0.685) |
| Reflections used in refinement | 15515 (1499) |
| Reflections used for R-free | 1264 (121) |
| R-work | 0.209 (0.688) |
| R-free | 0.221 (0.726) |
| Total non-hydrogen atoms | 4342 |
| macromolecules | 4279 |
| ligands | 56 |
| water | 7 |
| RMS(bonds) | 0.015 |
| RMS(angles) | 0.49 |
| Clashscore | 4.01 |
| Average B-factor | 69.85 |
| macromolecules | 69.17 |
| ligands | 93.63 |
| solvent | 66.69 |

**Figure S1|.**
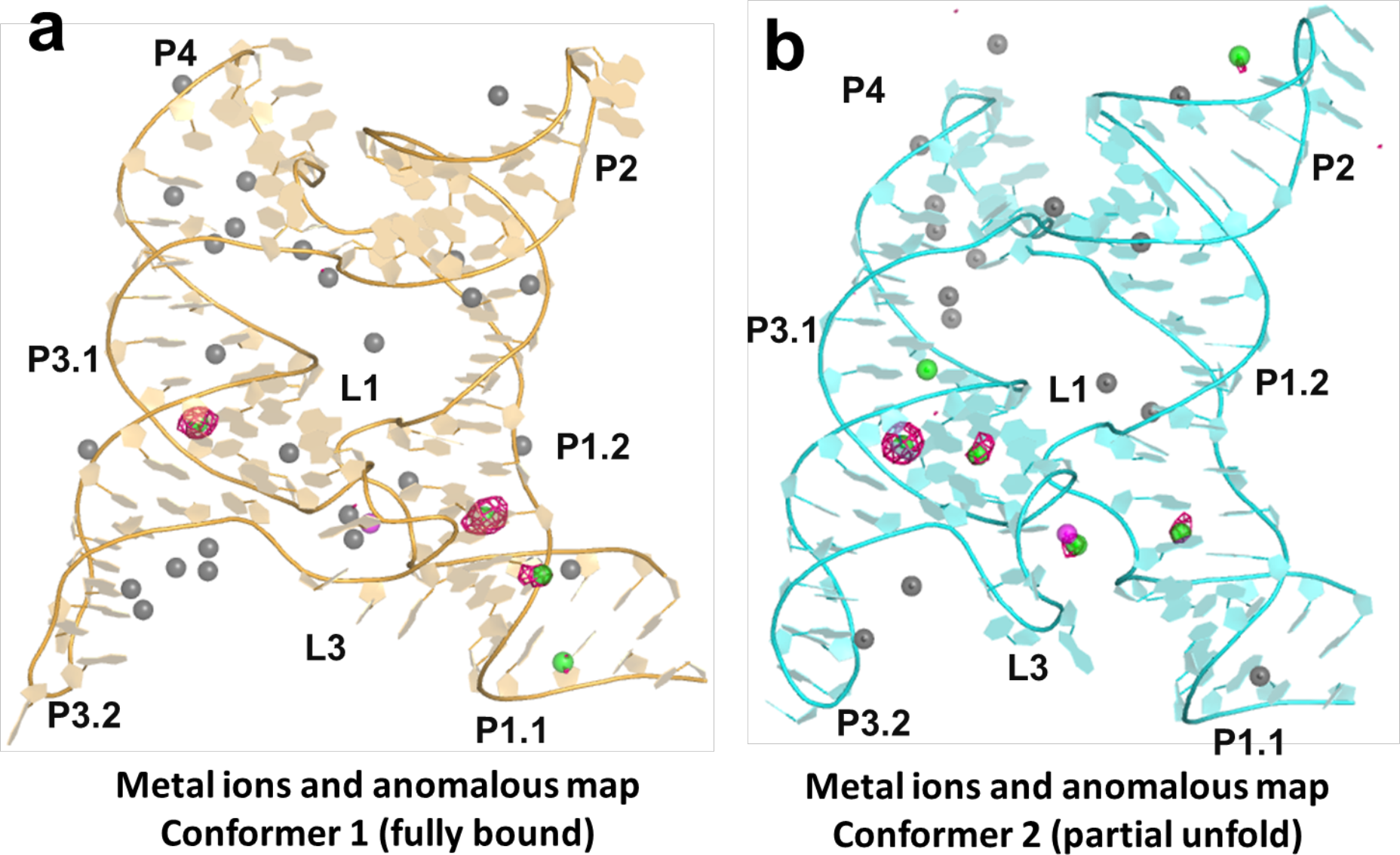
Overall ions and anomalous sites for the *X. oryzae yybP-ykoY* riboswitch crystal construct. **a)** Conformer 1 (orange) and **b)** Conformer 2 (cyan). Total ions within 5 Å of A chain include 2 Mn^2+^ (magenta), 45 Mg^2+^ (or Mn^2+^) (black), and 9 Sr^2+^ (green). The anomalous difference map, collected at 0.769 Å, is shown in pink mesh at level 4 σ. Placement of Sr^2+^ ions was determined by anomalous map and/or high electron density. Since Mn^2+^ has minimal anomalous signal at this wavelength and it is similar in ionic radius to Mg^2+^, it could not be differentiated from Mg^2+^. Thus, Mn^2+^ (2.5 mM in the crystal) could partially or fully occupy sites denoted as Mg^2+^ (30 mM in the crystal). Binding site Mn^2+^ were predicted to be so based on previous analysis of the *L. lactis* structure, but were not confirmed by this structure.

**Figure S2.**
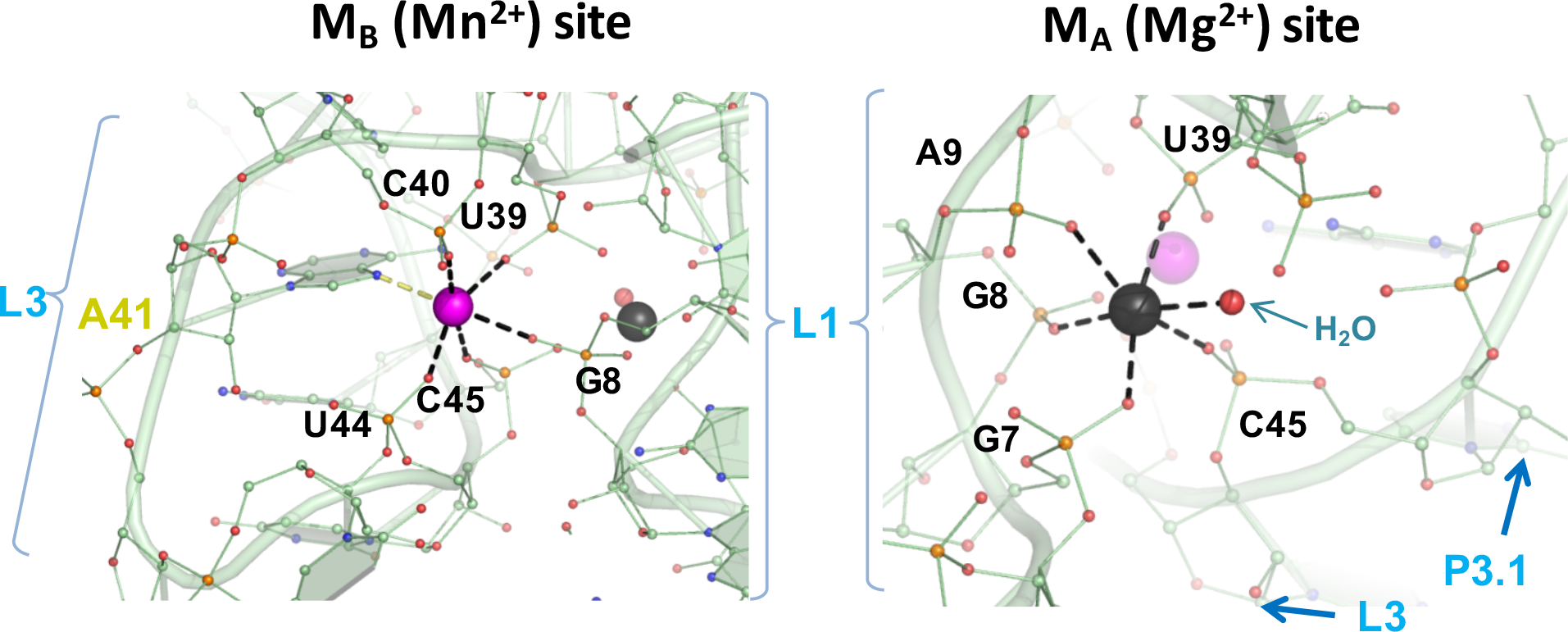
Metal binding sites in the *L. lactis yybP-ykoY* structure. Close-up view of the M_B,Mn_ (purple) and M_A,Mg_ (black) metal binding sites showing different contacts with surrounding L3 and L1 residues. The oxygen atoms of water molecules are shown as red spheres.

**Figure S3.**
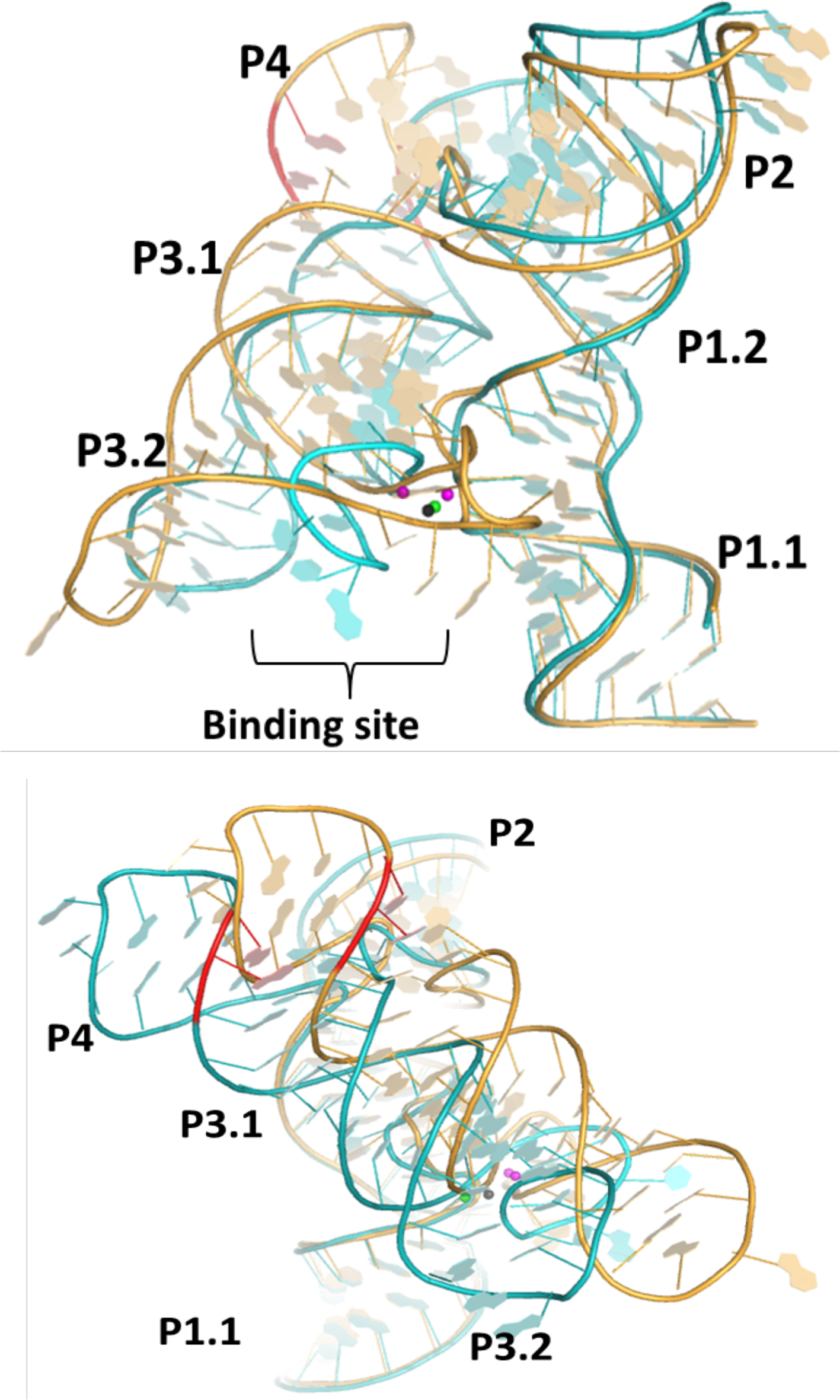
Overall comparison of Conformers 1 and 2. When aligned by L1, there is also a shift of the P3/P4 helical stack between Conformers 1 (orange) and 2 (cyan). The *L. lactis* structure (not shown) is intermediate but more similar to Conformer 1 when aligned in this way.

### Comparison of Xory Mn^2+^ bound structures to Cd^2+^ bound structures

With respect to the recent finding that Cd^2+^ can also bind to the *yybP-ykoY* riboswitch^1^, our smFRET results and unpublished data from a yybP-ykoY and Broccoli-based fluorescent sensor agree with this conclusion. However, our structure (at 2.85 A resolution) and previous ones with Mn^2+^ cannot provide strong evidence for or against their intriguing argument for heptacoordination as the mechanism of specificity for Mn^2+^. While Cd^2+^ is heptacoordinate in their Llac-MntP structure (PDB ID 6CC3), with water as its seventh ligand, it is not clear how that water could enable a mechanism of specificity for Mn^2+^ against other metals. Further, in the highest-resolution structure (6CB3), Cd^2+^ at M_B_ was found to be hexacoordinate. The Llac-alx structure (6CC1) is also modelled as bound in a heptacoordinate fashion to M_B_, with the seventh ligand here coming from a second phosphoryl oxygen from the same phosphate of U44. However, this structure is similar to all of our Mn^2+^-bound structures, in that the resolution is not high enough to show the subtle difference in orientation of a phosphate that would be required to distinguish hexa-from hepta-coordination. As for the M_A_ site, our Cd^2+^-only smFRET data suggest that M_A_ probably prefers either Mg or Mn over Cd^2+^. Upon inspection of their data, M_A_ does not appear clearly heptacoordinate in any of their structures. It is actually octacoordinate in the Llac with Cd/Mg/Ba (PDB 6CB3) and another claim (Llac-MntP) relies on placed waters not seen in the electron density. A very high-resolution structure of a *yybP-ykoY* riboswitch with Mn^2+^ is required to address this interesting question. In any case, taken together with our structures, these agree with the finding of flexibility in the Mn-binding area, even when the riboswitch ligand is bound. It is a separate biological argument whether Cd^2+^ is a relevant ligand, as it in the past was often considered toxic and xenobiotic for most organisms, and its high-affinity binding to enzymes at sites of other metals is generally considered aberrant^2^.

### Global structural dynamics of docked structures of aptamers in the MD simulations

To elucidate structural dynamics of Mn^2+^ sensing riboswitch, we performed a set of explicit solvent MD simulations on microseconds time scale. In addition to simulations containing both M_A_ and M_B_ ion binding sites occupied by divalent ions, we performed a set of simulations testing the effect of replacement of these divalent ions by monovalents in each ion binding position separately or in both of them simultaneously (see **Table S2**).

The global structural dynamics of the studied systems was visualized by the B-factors calculated per residues (**Figure S4**). All simulations revealed similar trends in global structural dynamics except of behavior of L3 loop, which was found to be sensitive to type of ions in the M_A_ and M_B_ ion binding sites (see below). All stems exhibited high structural stability in all simulations. We observed larger fluctuations on the tips of P2, P3, and P4 stems. These fluctuations were found to be caused by unstacking of the L2 nucleotide, which should be considered as inherent part of native structural dynamics of GNRA tetraloops^3–7^. In the case of simulations of *L. lactis* structure, B-factor values indicated also large fluctuations of the P2 stem. Detailed analyses revealed that these fluctuations were connected with a reversible bending of the P2 stem and its movement toward and away from the P4 stem. This global movement was correlated with flipping of ε torsion angle of A28 nucleotide, which fluctuated between two states (see **Figure S5**).

**Figure S4.**
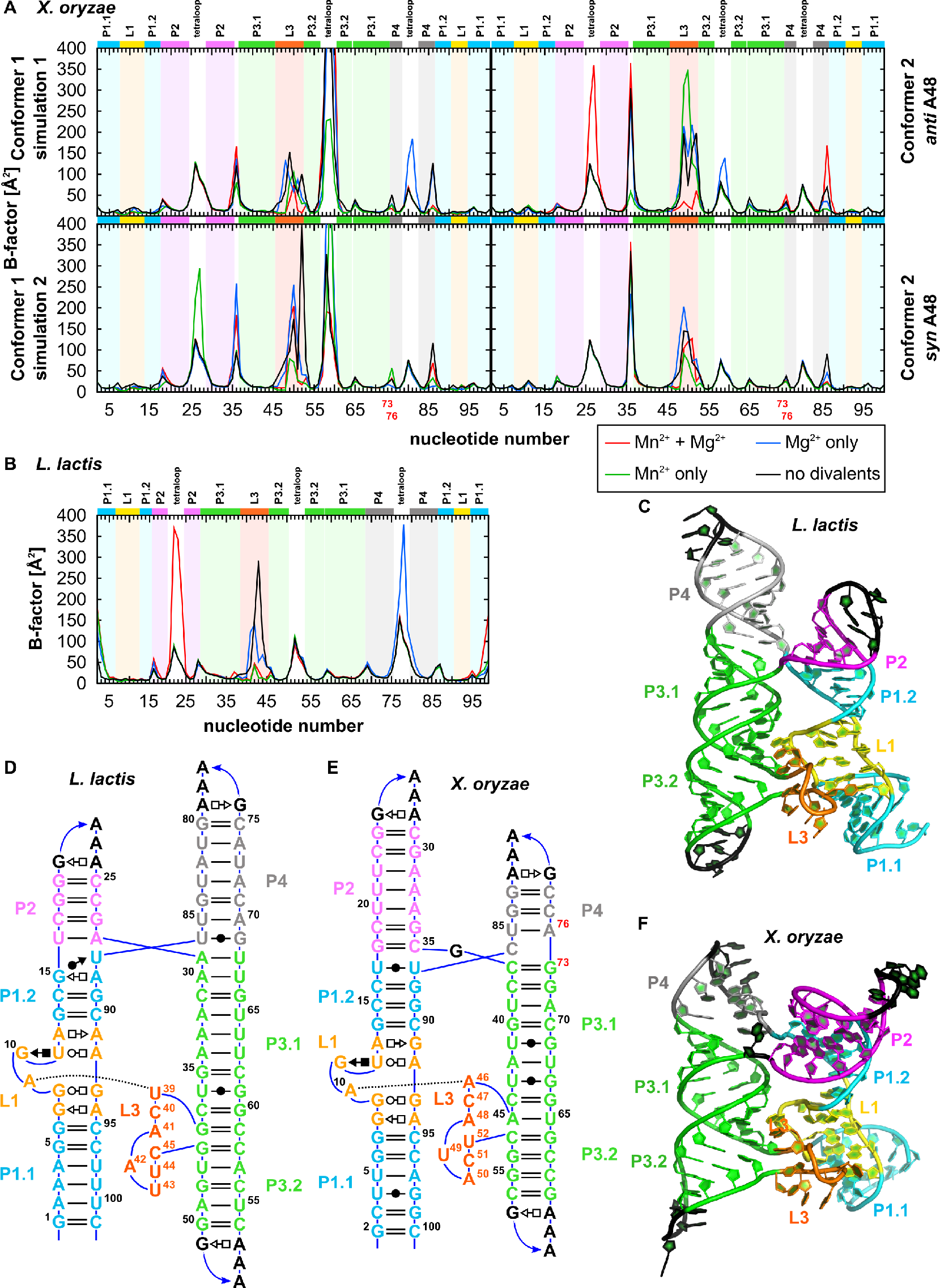
The average mass-weighted B-factor as a function of residue number obtained in MD simulations of Mn^2+^ sensing riboswitch from A) *Xanthomonas oryzae* or B) *Lactococcus lactis* (see Table 1 in the main text for full list of the simulations). The colors of background bars correspond to the regions in the secondary and 3D structures shown on panels C-F.

**Figure S5.**
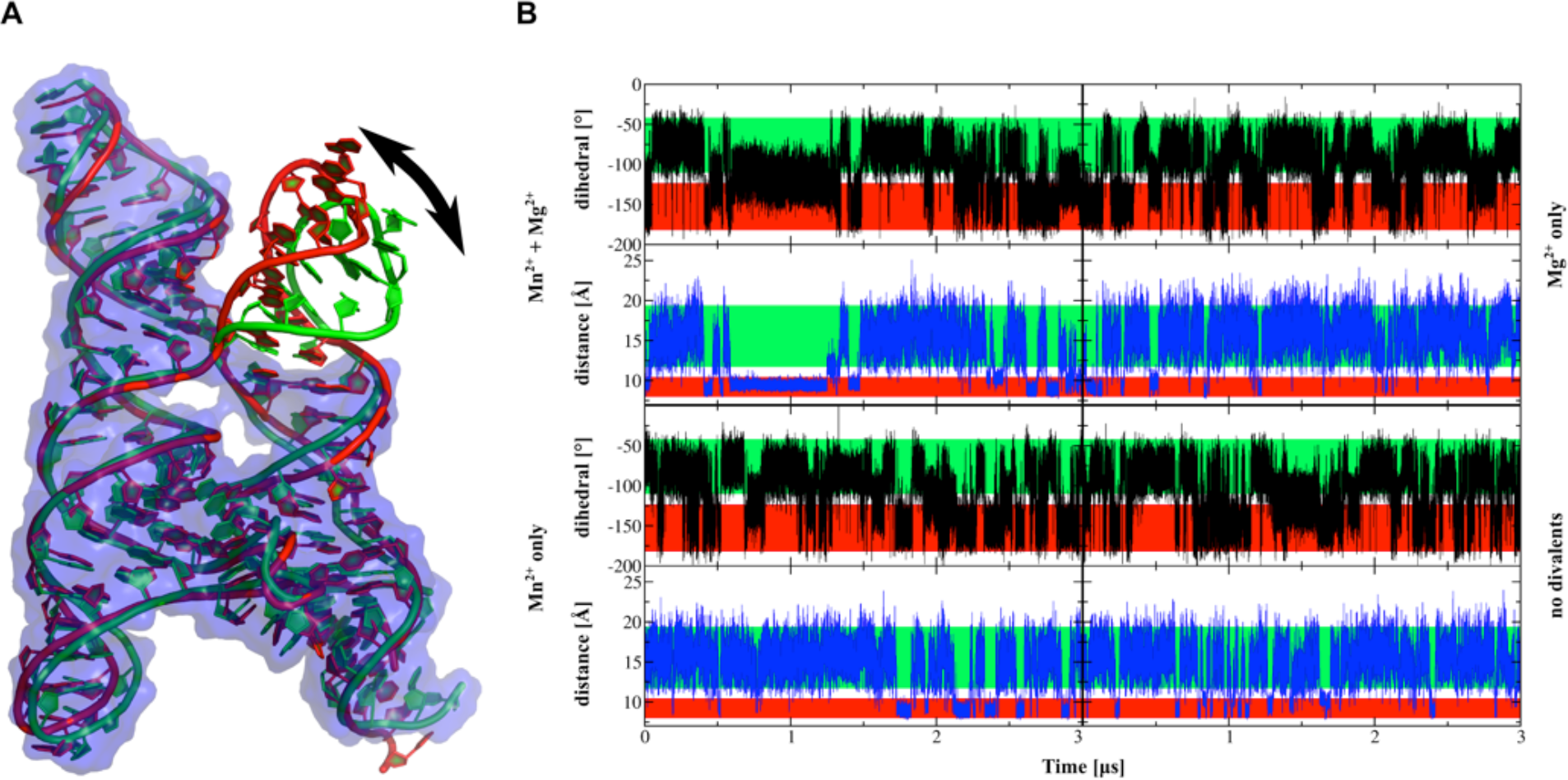
A) Superposition of the open and closed conformation (in green and red, respectively) observed in MD simulation of the aptamer from *L. lactis*. B) Time dependence of ε dihedral angle of A28 (black line) and distance between P2 (center of mass of C17-G20 backbone) and P4 (center of mass of C71-A72 backbone) stems (blue line). The green and the red strips on the background correspond to the open (native crystal-like) and closed conformation, respectively.

### MD simulations confirmed syn orientation of A48 in L3 loop of X. oryzae structure Conformer 2

The electron density of the conformer 2 of the *X. oryzae* structure revealed adenine A48 in *syn-* rather than *anti*-conformation. In order to verify this unusual glycosidic bond orientation within the given structural context, we used two different refinements of the conformer 2 structure (**Figure S6**) as starting structures for subsequent MD simulations. These simulations aimed to indirectly probe the orientation of A48 glycosidic bond via analyses of compatibility of a given orientation of A48 with its structural environment reflecting the native A48 orientation through molecular interactions. We thus expected that while simulation starting from native orientation of A48 will fluctuate around starting structure conformation, the non-native orientation of A48 in the starting structure should result in structural changes in initial phase of MD simulation. For sake of completeness, it is also possible that both orientations of A48 might coexist in the crystal lattice ensemble and electron density represents ensemble-averaged picture of A48, which is structurally compatible with its structural environment in both orientations. In such case, both A48 orientations would be equally tolerated by their structural environment and thus both would be equally stable in MD simulations.

Therefore, we performed MD simulations of conformer 2 both with *syn*- and *anti*-oriented A48. Namely, we compared simulations having their ion binding sites occupied by corresponding native divalent ions, i.e., by Mg^2+^ and Mn^2+^ in M_A_ and M_B_, respectively (see **Table S2**). We observed that the simulation started from structure with *syn*-oriented A48 stably fluctuated around crystal conformation during the entire 2 μs simulation. In contrast, the simulation started with *anti*-oriented A48 revealed significant structural changes during initial phase of the simulation. Namely, the A48 nucleobase was shifted already in the initial geometrical optimization, so that it was coordinated to the Mn^2+^ ion by N7 nitrogen, while N6 exocyclic amino group was repelled away from the Mn^2+^ ion. This movement was accompanied by reconformation of the sugar-phosphate backbone between U52 and C53 that shifted away from A48 to avoid sterical clash (**Figure S7**). Subsequently, in the early stages of the simulations, namely during thermal equilibration, such reconformation of U52-C53 sugar-phosphate backbone resulted in destabilization of the U52(H3)…A48(O2’) hydrogen bond and exposure of the U52 into solvent (**Figure S7**). All these rapid structural changes should not be considered as native structural dynamics of the riboswitch and are unambiguously a consequence of starting structure bias, namely non-native refinement of A48 *anti* orientation. Thus, in agreement with the observed electron density, we found that *anti*-oriented A48 is not compatible with the overall structure of Conformer 2 containing open-conformation of loop L3. Thus, crystallographic data together with MD simulations provide clear evidence of *syn*-orientated A48 in this particular L3 loop conformation; the MD simulations of Conformer 2 started with *anti*-A48 were not further analyzed and discussed.

For the sake of completeness, it is worth noting that MD simulations of Conformer 1 (containing closed conformation of loop L3 and A48 clearly resolved as *anti*-orientated) did not reveal any rapid structural changes during the early stage relaxation that would point to any doubts of the refinement of the L3 loop conformation.

**Figure S6.**
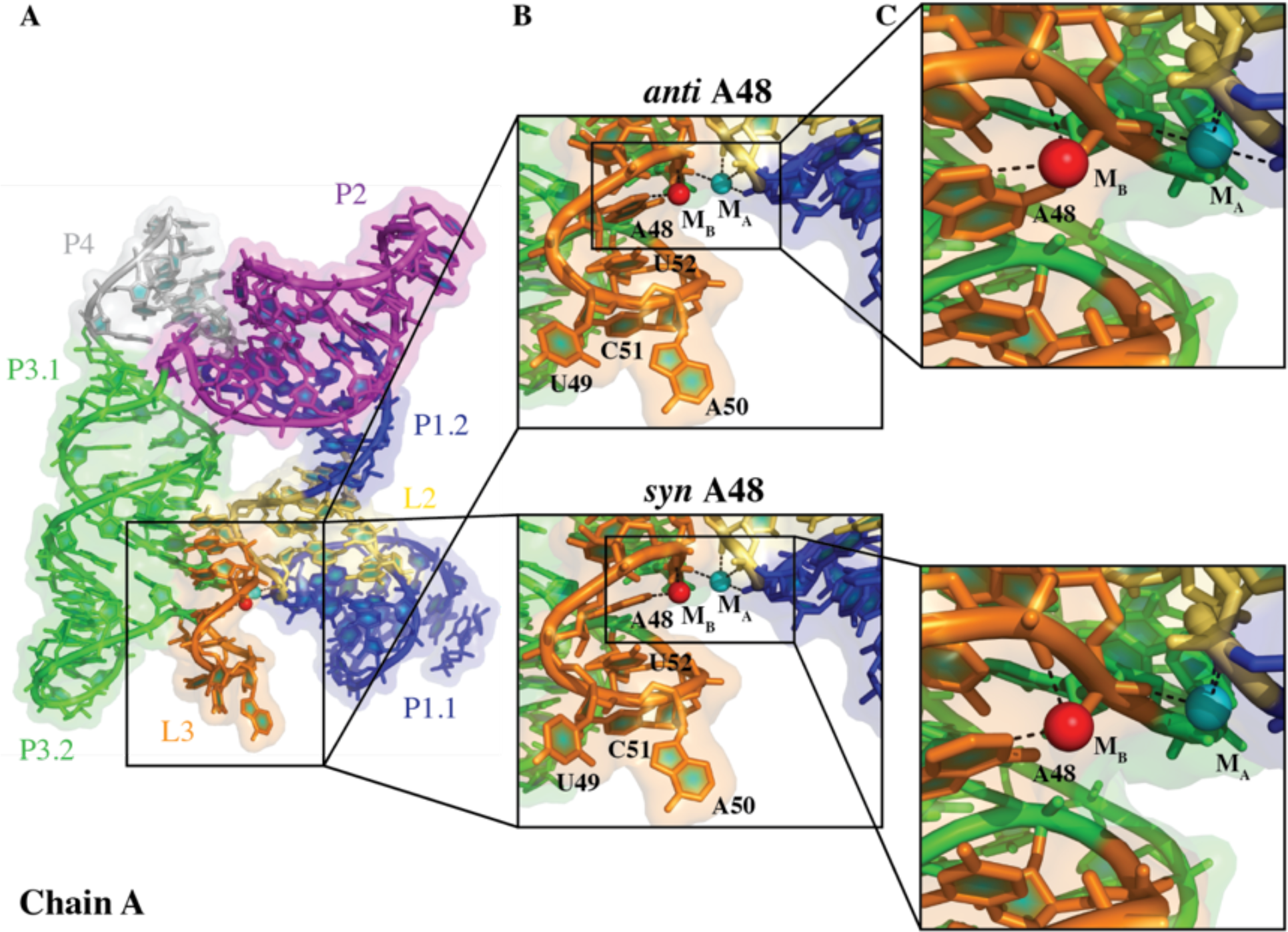
Structure of *X. oryzae* aptamer domain of conformer 2 showing alternative *syn* and *anti* orientations of A48.

**Figure S7.**
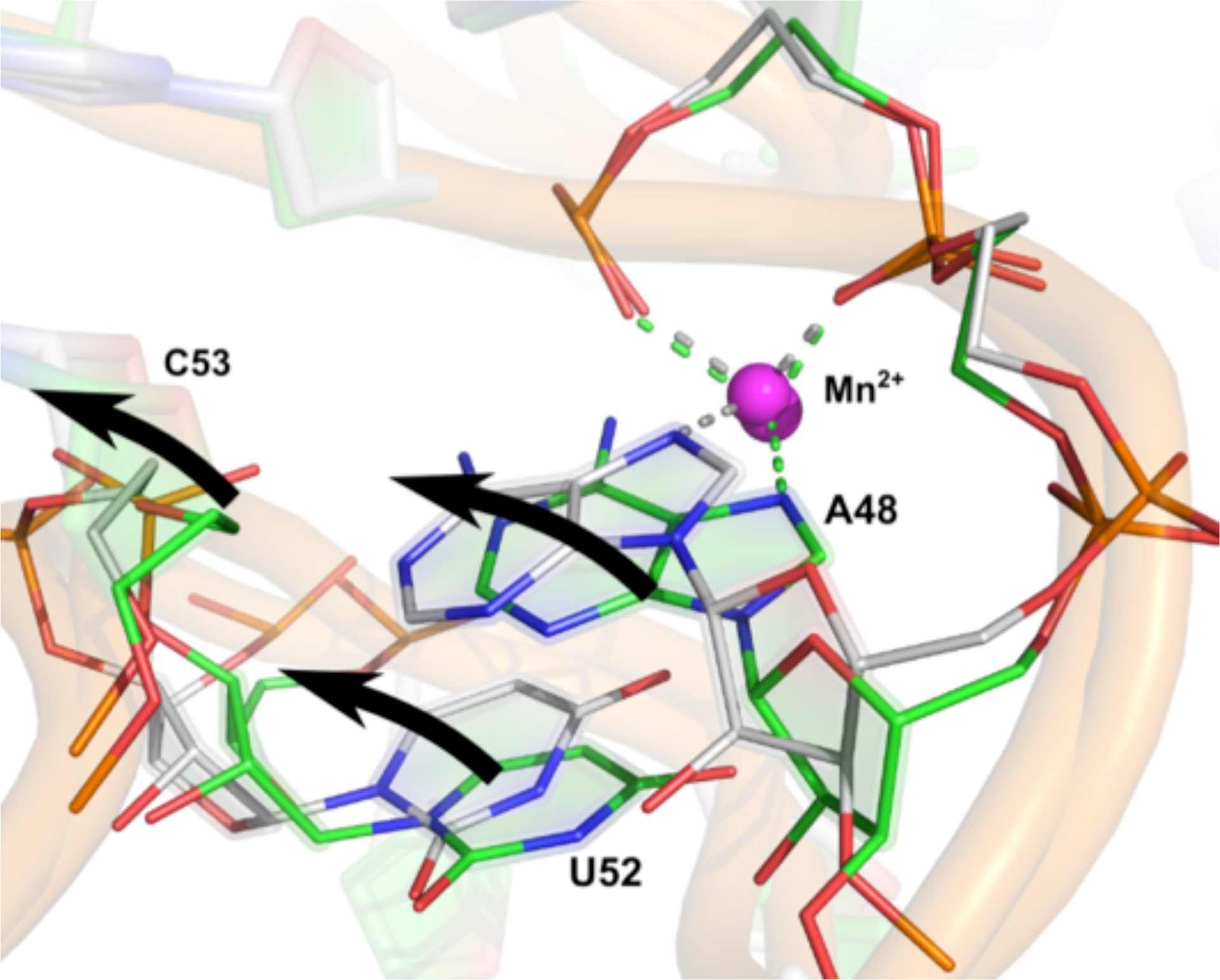
Structural changes observed during early stages of the MD simulations of *X. oryzae* conformer 2 with *anti*-orientation of A48. Green structure corresponds to crystal structure, while the silver refers to the structure after initial rearrangements during these early stages.

**Table S2 -.** Percentage of native stacking preserved in L3 loop during MD simulations (based on annotations calculated by baRNAba software) and percentage of occupancy of ions in the ion binding sites.

| Ions |  | Stacking interactions<br>[%] |  |  |  |  |  | Occupancy of M<sub>A</sub><br>[%] |  |  |  |  | Occupancy of M<sub>B</sub><br>[%] |  |  |  |  |  |
| --- | --- | --- | --- | --- | --- | --- | --- | --- | --- | --- | --- | --- | --- | --- | --- | --- | --- | --- |
| M<sub>A</sub> | M<sub>B</sub> | A9 | U39 | C40 | A41 | U44 | A42 | G7 | G8 | A9 | U39 | C45 | G8 | U39 | C40 | A41 | U44 | C45 |
|  |  | U39 | C40 | A41 | C45 | C45 | U44 | OP1 | OP2 | OP1 | OP1 | OP1 | OP1 | OP2 | OP2 | N7 | OP1 | OP2 |
| *L.lactis* | Mg<sup>2+</sup> Mn<sup>2+</sup> | 100 | 98 | 77 | 98 | 92 | 76 | 100 | 100 | 100 | 100 | 100 | 100 | 100 | 100 | 100 | 100 | 100 |
|  | K<sup>+</sup> Mn<sup>2+</sup> | 99 | 99 | 78 | 96 | 65 | 83 | 100 | 100 | 100 | 100 | 2 | 100 | 100 | 100 | 100 | 100 | 100 |
|  | Mg<sup>2+</sup> K<sup>+</sup> | 99 | 99 | 2 | 4 | 5 | 6 | 100 | 100 | 100 | 100 | 100 | 100 | 100 | 98 | 6 | 0 | 100 |
|  | K<sup>+</sup> K<sup>+</sup> | 100 | 93 | 64 | 88 | 36 | 42 | 95 | 98 | 95 | 100 | 0 | 27 | 99 | 78 | 97 | 42 | 67 |
|  |  | A10 | A46 | C47 | A48 | C51 | A50 | G8 | G9 | A10 | A46 | U52 | G9 | A46 | C47 | A48 | C51 | U52 |
|  |  | A46 | C47 | A48 | U52 | U52 | C51 | OP2 | OP2 | OP1 | OP1 | OP1 | OP1 | OP2 | OP2 | N7/1 | OP1 | OP2 |
| *X.ory*_cf2_syn | Mg<sup>2+</sup> Mn<sup>2+</sup> | 100 | 100 | 100 | 66 | 61 | 63 | 100 | 100 | 100 | 100 | 0 | 0 | 100 | 100 | 100 | 0 | 0 |
|  | K<sup>+</sup> Mn<sup>2+</sup> | 91 | 100 | 100 | 81 | 86 | 89 | 99 | 99 | 100 | 100 | 0 | 0 | 100 | 100 | 100 | 0 | 0 |
|  | Mg<sup>2+</sup> K<sup>+</sup> | 100 | 99 | 87 | 69 | 82 | 89 | 100 | 100 | 100 | 100 | 0 | 65 | 68 | 67 | 68 | 17 | 17 |
|  | K<sup>+</sup> K<sup>+</sup> | 96 | 99 | 89 | 70 | 90 | 67 | 95 | 98 | 98 | 98 | 45 | 6 | 90 | 67 | 68 | 10 | 67 |
| *X.ory*_cf1_1 | Mg<sup>2+</sup> Mn<sup>2+</sup> | 100 | 100 | 54 | 92 | 84 | 11 | 0 | 100 | 0 | 100 | 100 | 100 | 100 | 100 | 100 | 100 | 100 |
|  | K<sup>+</sup> Mn<sup>2+</sup> | 98 | 96 | 70 | 75 | 70 | 47 | 99 | 100 | 100 | 100 | 1 | 100 | 100 | 100 | 100 | 100 | 100 |
|  | Mg<sup>2+</sup> K<sup>+</sup> | 99 | 86 | 5 | 1 | 5 | 2 | 0 | 100 | 0 | 100 | 100 | 25 | 24 | 21 | 6 | 5 | 24 |
|  | K<sup>+</sup> K<sup>+</sup> | 97 | 99 | 76 | 69 | 78 | 86 | 79 | 85 | 79 | 92 | 11 | 20 | 85 | 74 | 82 | 19 | 38 |
| *X.ory*_cf1_2 | Mg<sup>2+</sup> Mn<sup>2+</sup> | 100 | 99 | 57 | 90 | 86 | 3 | 0 | 100 | 0 | 100 | 100 | 100 | 100 | 100 | 100 | 100 | 100 |
|  | K<sup>+</sup> Mn<sup>2+</sup> | 93 | 97 | 82 | 45 | 86 | 11 | 99 | 100 | 99 | 100 | 1 | 100 | 100 | 100 | 100 | 100 | 100 |
|  | Mg<sup>2+</sup> K<sup>+</sup> | 100 | 99 | 84 | 2 | 15 | 24 | 0 | 100 | 0 | 100 | 100 | 99 | 99 | 100 | 25 | 4 | 100 |
|  | K<sup>+</sup> K<sup>+</sup> | 87 | 95 | 54 | 37 | 47 | 60 | 35 | 94 | 62 | 97 | 5 | 72 | 34 | 83 | 76 | 59 | 23 |

**Figure S8.**
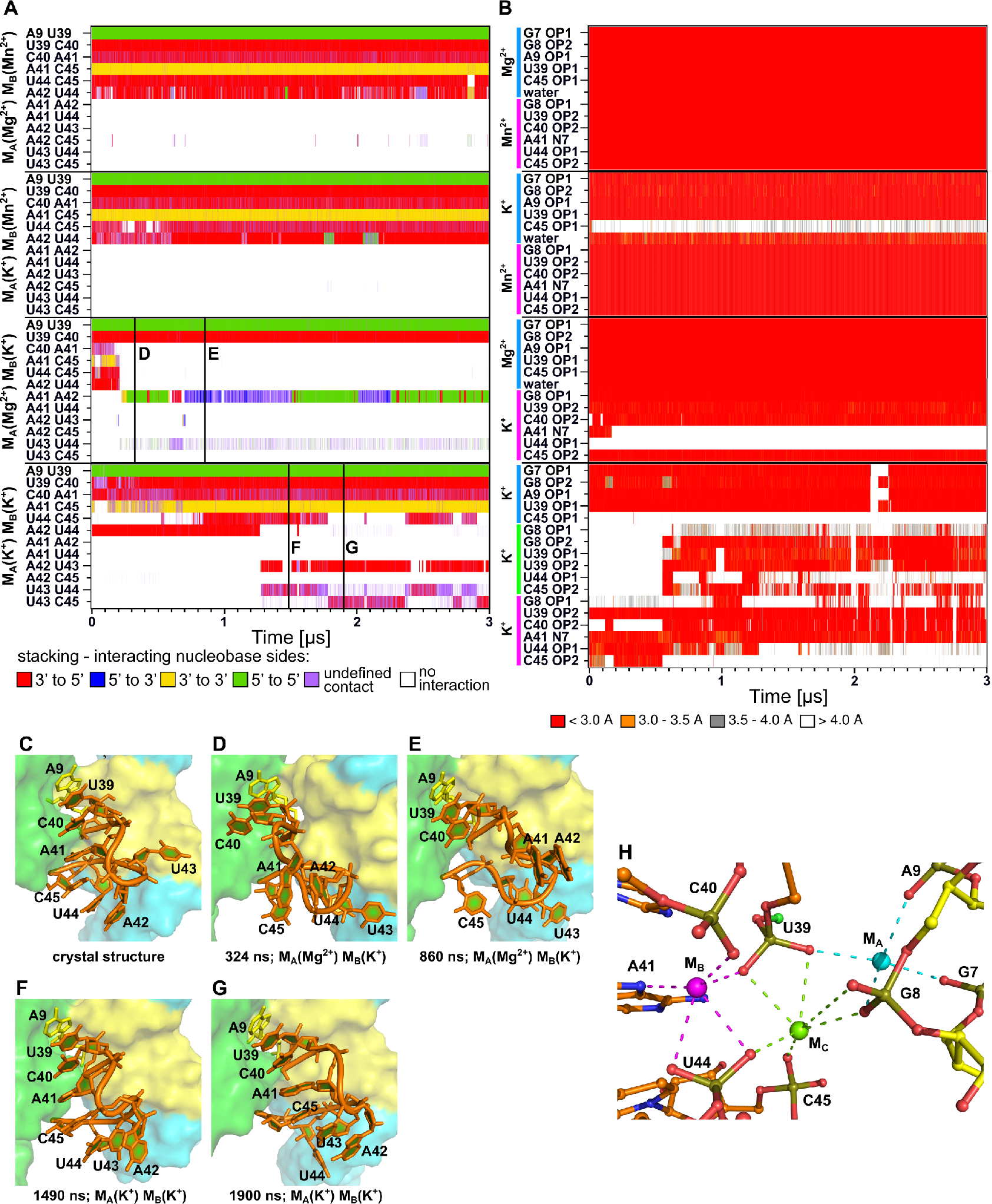
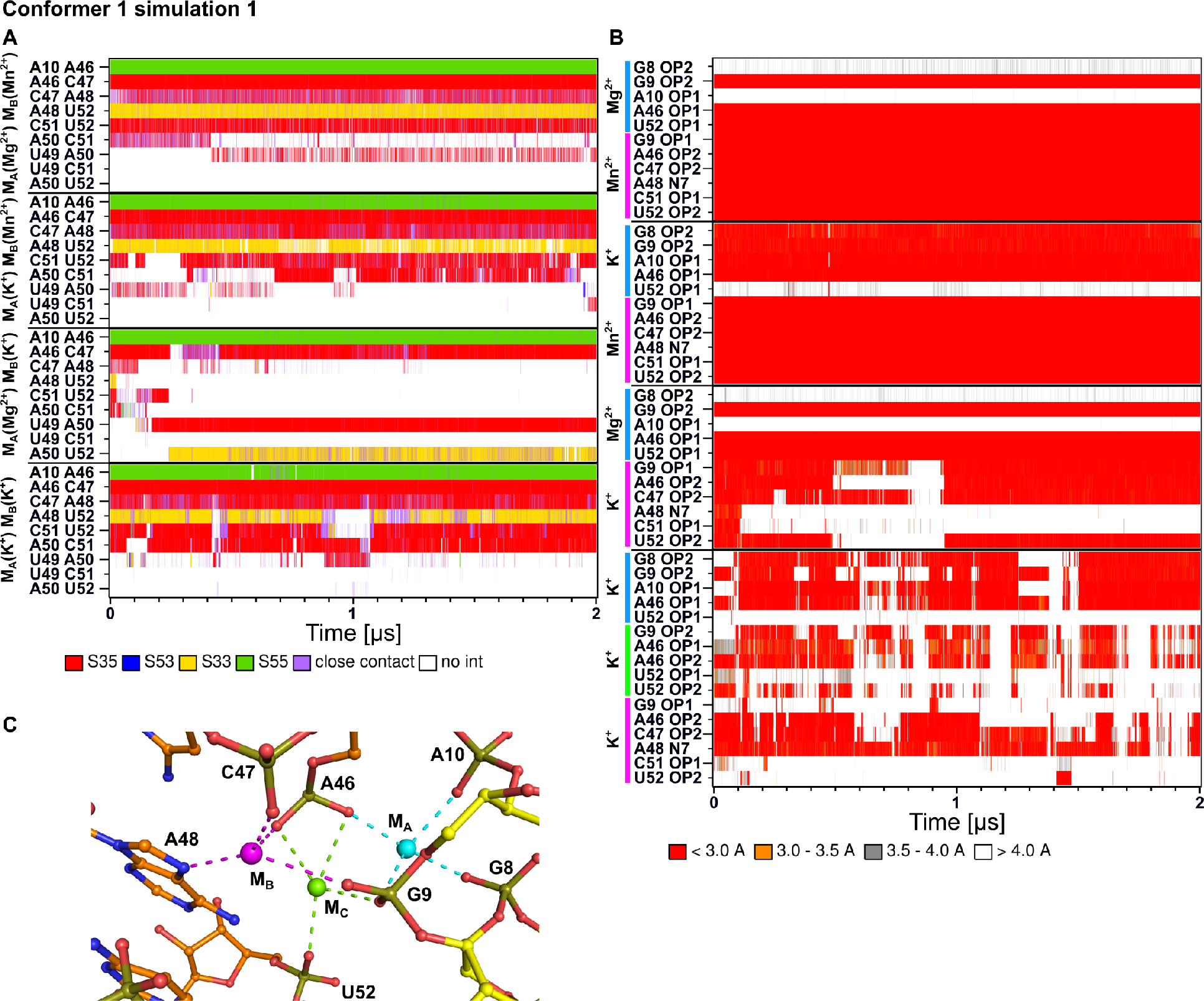
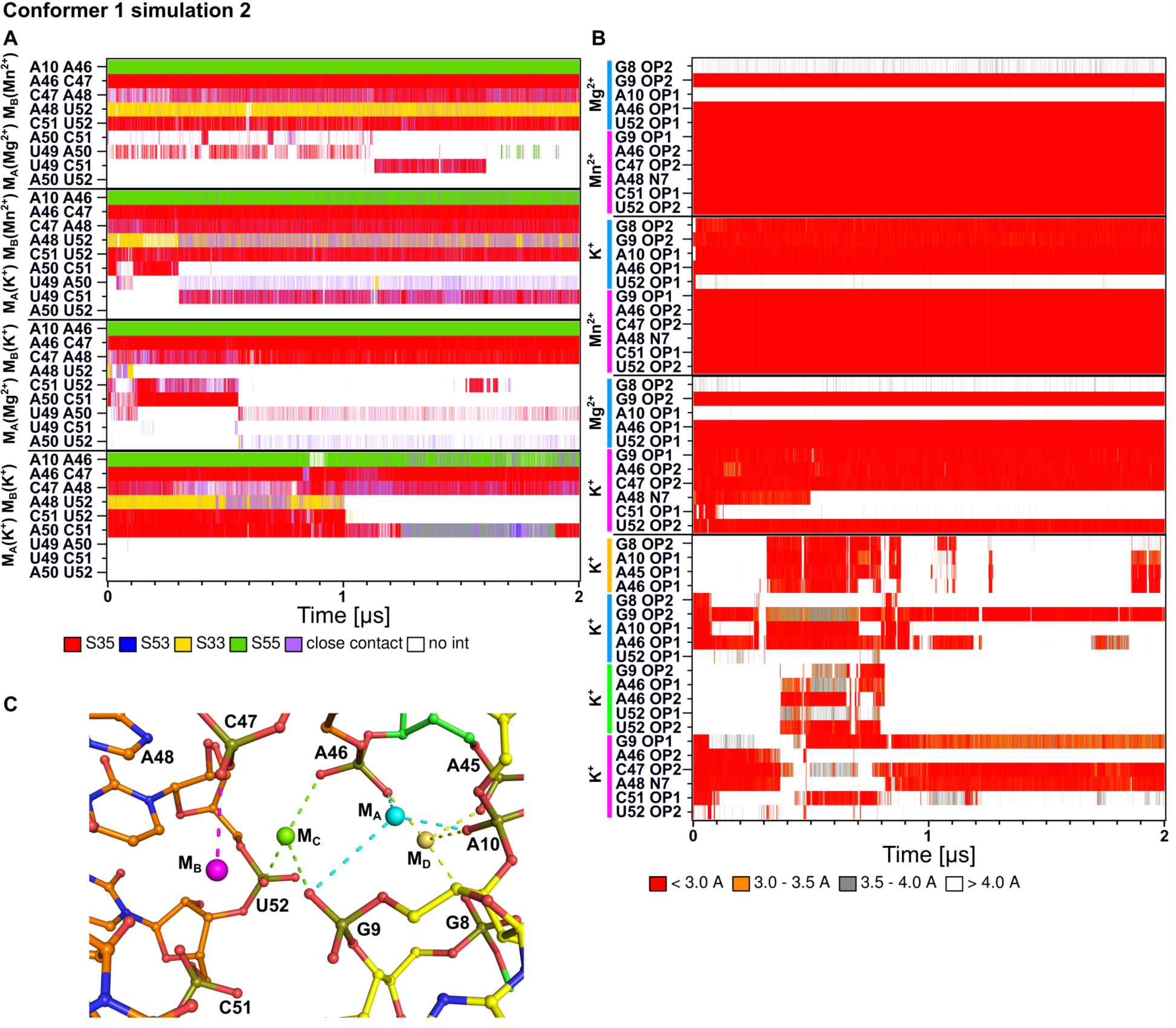
Conformational behavior of the L3 loop and MA and MB ion binding sites in MD simulation of *L. lactis* aptamer. A) Time evolution of the stacking pattern of the loop; the colors correspond to different mutual orientations of nucleobases in stacking interactions indicated by the corresponding faces (3’-face and 5’-face) exposed to the stacking interaction by stacked nucleobases. B) Time evolution of ligand-ion interactions in the ion binding sites. C-G) Close view on stacking pattern, panel C corresponds to crystal structure, while the others depict structures observed in MD simulations. H) New ion binding sites in simulation where both divalents were replaced by K^+^.

**Figure S9.**
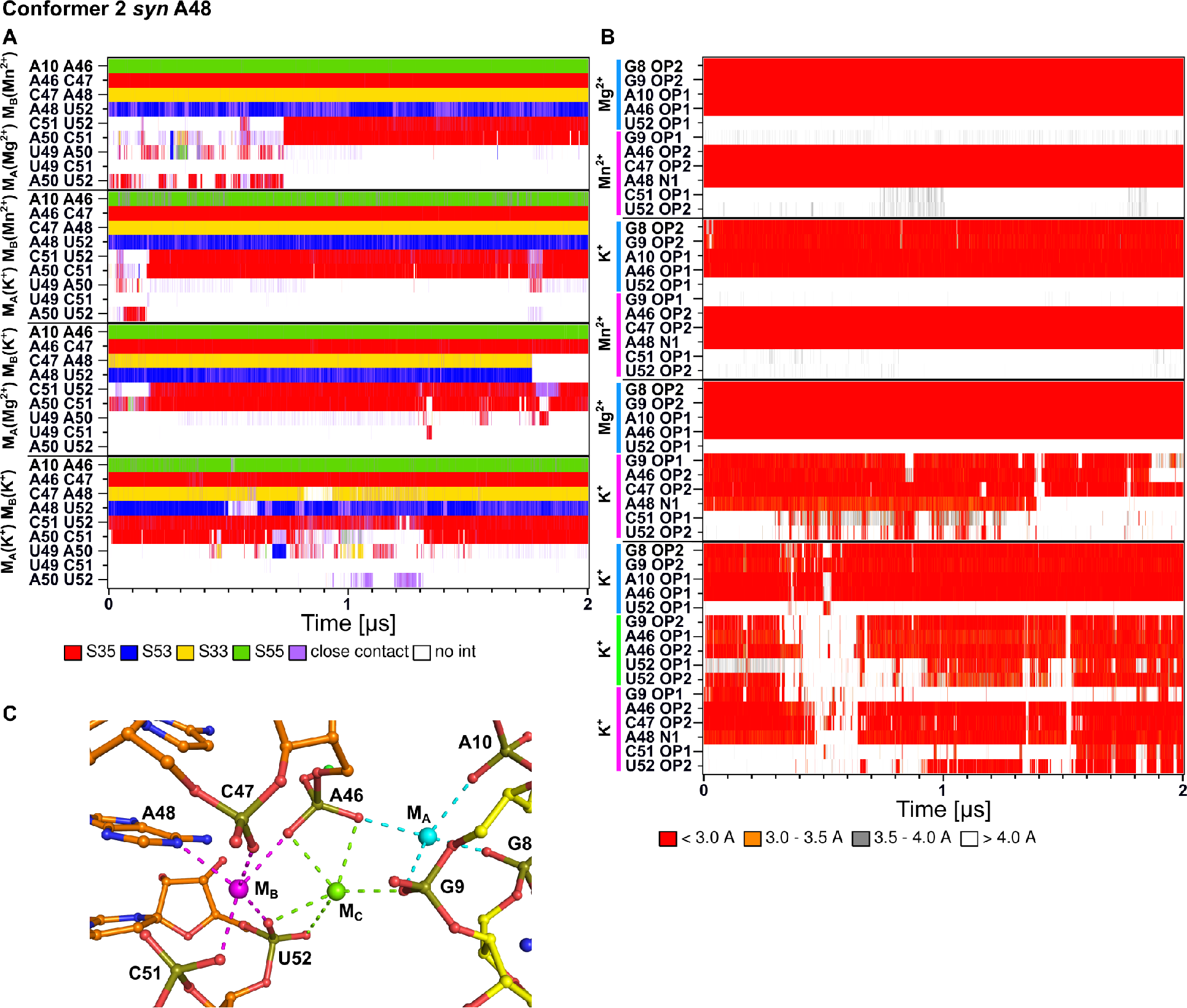
Conformational behavior of the L3 loop and MA and MB ion binding sites in MD simulation of *X. oryzae* aptamer. **A)** Time evolution of the stacking pattern. **B)** Time evolution of ligand-ion interactions in the ion binding sites. **C)** New ion binding sites in simulation where both divalents were replaced by K^+^.

### Stacking pattern of L3

Most of the nucleobases from L3 loop form continuous stacking pattern (**Figure S4 D,E**, Figure S11. This segment is further stacked by its 5’-end on adenine A10 (A9 according to numbering of *L. lactis*) from L1 loop, which forms type I A-minor interaction with G66=C44 (G61=C37 according to numbering of *L. lactis*) base pair. This A-minor interaction together with part of the stacking pattern formed by the A10(A9) adenine and two nucleotides at 5’-end of L3 loop (i.e., A10|A46|C47 and A9|U39|C40 in simulation of structure from *X. oryzae* and *L. lactis*, respectively) represent rather rigid part of the stacking pattern, which was found to be stable in all ionic conditions (**Figures S8**–**S9** and **Table S2**). In addition to the A-minor interaction making tertiary contact between L1 and L3 loops, the above-mentioned A10|A46|C47 (A9|U39|C40) part of the stacking pattern is stabilized also by other tertiary interactions such as hydrogen bonding of C47 (C40 according to numbering of *L. lactis*) cytosine with riboses of C53 and A45 (G46 and G38 in *L. lactis*) of the P3.2 stem (see Figures S11 and S12 for evolution of all tertiary contacts between A10|A46|C47 (A9|U39|C40) part of the stacking pattern and its structural environment). The insensitivity of this region to types of ions in M_A_ and M_B_ sites suggests that the L1-L3 loop tertiary contact mediated by this pattern might be formed in all ionic conditions even if M_A_ and M_B_ sites are not yet properly formed and occupied by the corresponding divalent ions.

The rest of stacking pattern of L3 loop (i.e., A48|U52|C51|A50 and A41|C45|U44|U43 in *X. oryzae* and *L. lactis* structures, respectively) showed different dynamics depending mostly on the type of ion in M_B_ site (**Figures S8–S9** and **Table S2**). The presence of Mn^2+^ ion in M_B_ significantly stabilized native stacking pattern in L3 loop, while this pattern was destabilized in simulations where the Mn^2+^ ion was replaced by K^+^. Surprisingly, when both divalent ions were replaced by K^+^ ions, the stability of L3 loop stacking pattern was higher than in case where only Mn^2+^ ion in M_B_ site was replaced by monovalent while M_A_ was occupied by Mg^2+^ ion, though still less stable than in simulations having also M_B_ site occupied by Mn^2+^. In order to explain this observation, we hypothesize that stacking pattern of L3 loop represents inherently quite stable conformation of this loop capable of remaining stable on at least microsecond time scale. However, when M_A_ binding site is occupied by Mg^2+^, unlike K^+^, it enforces proper positioning of A46 and U52 (U39 and C45 according *L. lactis* numbering) phosphates which leads to destabilization of the stacking pattern of L3. This stacking-structure-destabilization effect of Mg^2+^ can be overcompensated only by binding of Mn^2+^ ion into the M_B_ binding site.

**Figure S10.**
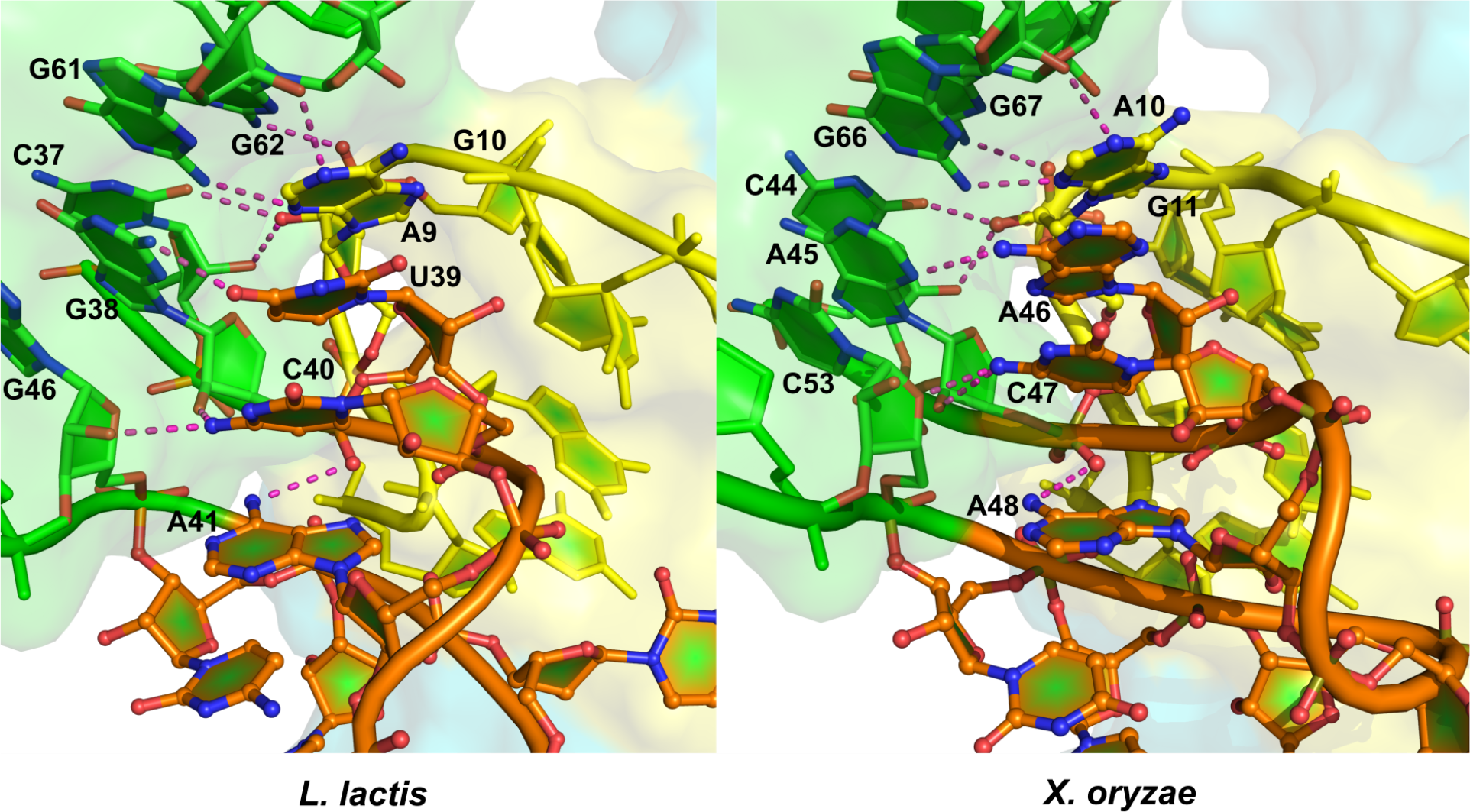
The stacking pattern formed by A10 (A9 according to numbering of *L. lactis*) and nucleobases of loop L3 with depicted tertiary interactions (see **Figure S11** for their structural stabilities in MD simulations).

**Figure S11.**
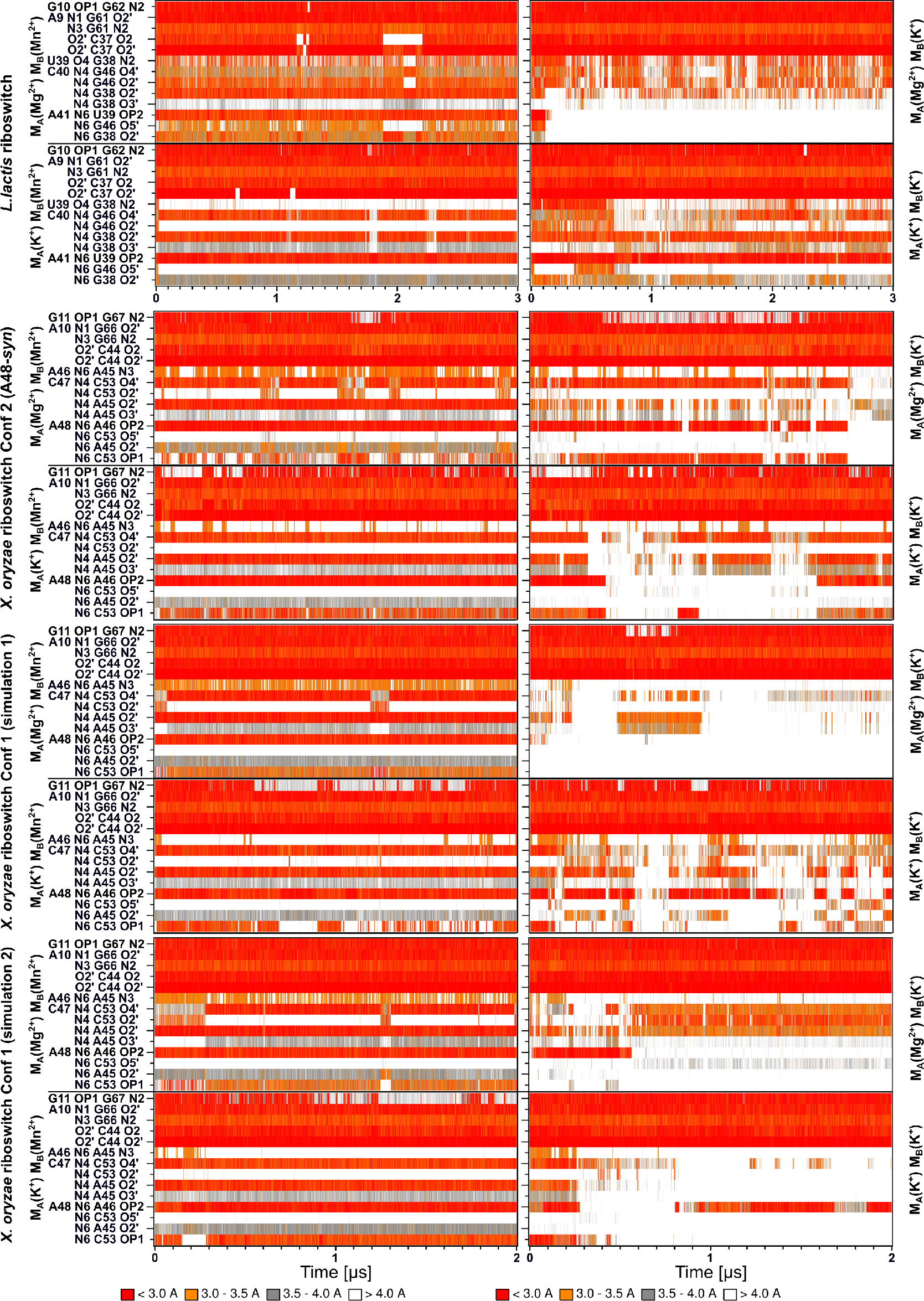
Time evolution of tertiary interactions of nucleotides included in the stacking pattern of L3 loop (see **Figure S10** for structural view of these tertiary contacts).

### Structural dynamics of SRL-like conformation of L1 in different structural contexts

Besides structures of whole aptamer which all remained in docked state in our simulations, we also performed simulations of the structures consisting only of P1.1, P1.2 and L1 (based on *X. oryzae* – Conformers 1 and 2 and *L. lactis* crystal structures, see Table S2). Such minimalistic structure obviously lacks all tertiary contacts to L3 loop and P3 stem and thus its dynamics should correspond to the dynamics of this particular part in undocked state.

The base-pairing (**Figure S12**) as well as backbone conformation were monitored and compared to the corresponding values observed in the simulations of complete aptamer, i.e., in docked state. Note that the overall conformation of L1 loop as well as base pairing in P1.1 stem (with exception of terminal AU pairs of the P1.1 stem in simulation of *L. lactis* aptamer which revealed base pair fraying) were entirely stable in the simulations of complete aptamers. The most significant structural changes were observed in simulation of the isolated P1.2|L1|P1.1 structure based on Conformer 2 of the *X. oryzae* structure. The disruption of G9(N1)…G93(N7) H-bond was followed by reconformation of G8-A94 from *t*SH into *c*WC base pairing and loss of the S-turn conformation of the sugar-phosphate backbone of G8-A10. Interestingly, the simulation of the same structural motif derived from the Conformer 1 of *X. oryzae* crystal structure showed less pronounced changes, in particular, the G9(N1)…G93(N7) H bond was broken and recreated several times. Finally, in case of the simulation based on *L. lactis* structure, the L1 loop resembled conformation observed in the complete aptamer; however, we observed rather significant unpairing in P1.1 stem. This may be due to a weaker P1.1 in *L. lactis* that is shorter by 1-bp and has three A-U base pairs capping the stem, as opposed to the two G-C base pairs in the *X. oryzae* structure.

In summary, the data indicate that when A10 is sequestered in the A-minor interaction, the P1.2|L1|P1.1 segment adopts a conformation resembling the topology of a canonical sarcin-ricin loop. Compared to the sarcin-ricin loop consensus sequence, the A10 nt is inserted between its GpU platform and the flexible region. In addition, the trans-Hoogsteen/Hoogsteen base pair in the flexible region adjacent to the GpU platform is replaced by a trans-Watson-Crick/Hoogsteen pair. When the A10 is not involved in the tertiary interaction, it instead destabilizes the SRL-like topology.

**Figure S12.**
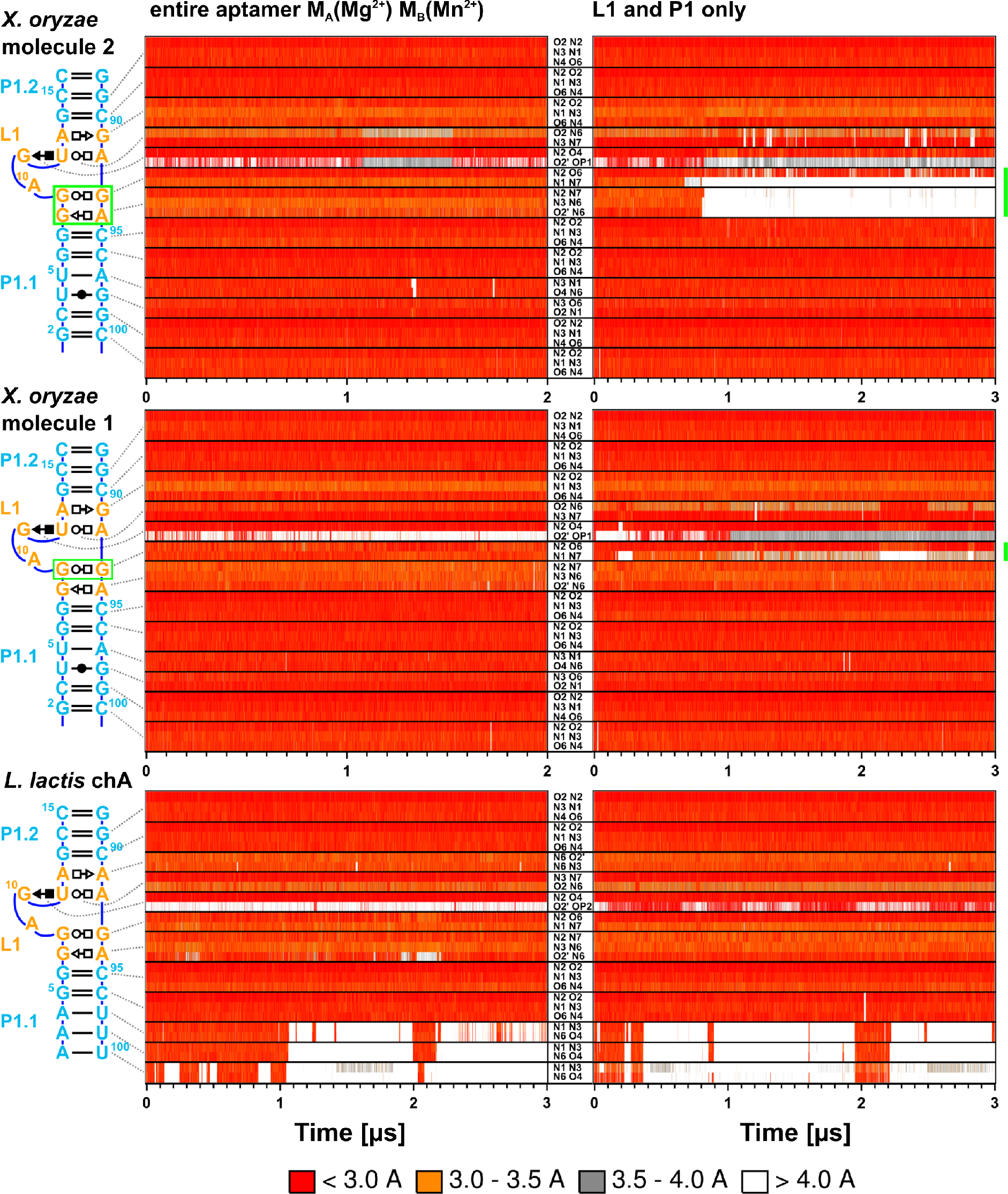
Time evolution of native base pairing interactions of SRL-like motif of L1 in entire aptamer and in model structure (consisting only of P1.1, P1.2 and L1 and thus lacking A-minor interaction of A10 in *X. oryzae* and A9 in *L. lactis*, to L3 loop and P3 stem). The most labile interactions are highlighted in green.

**Figure S13.**
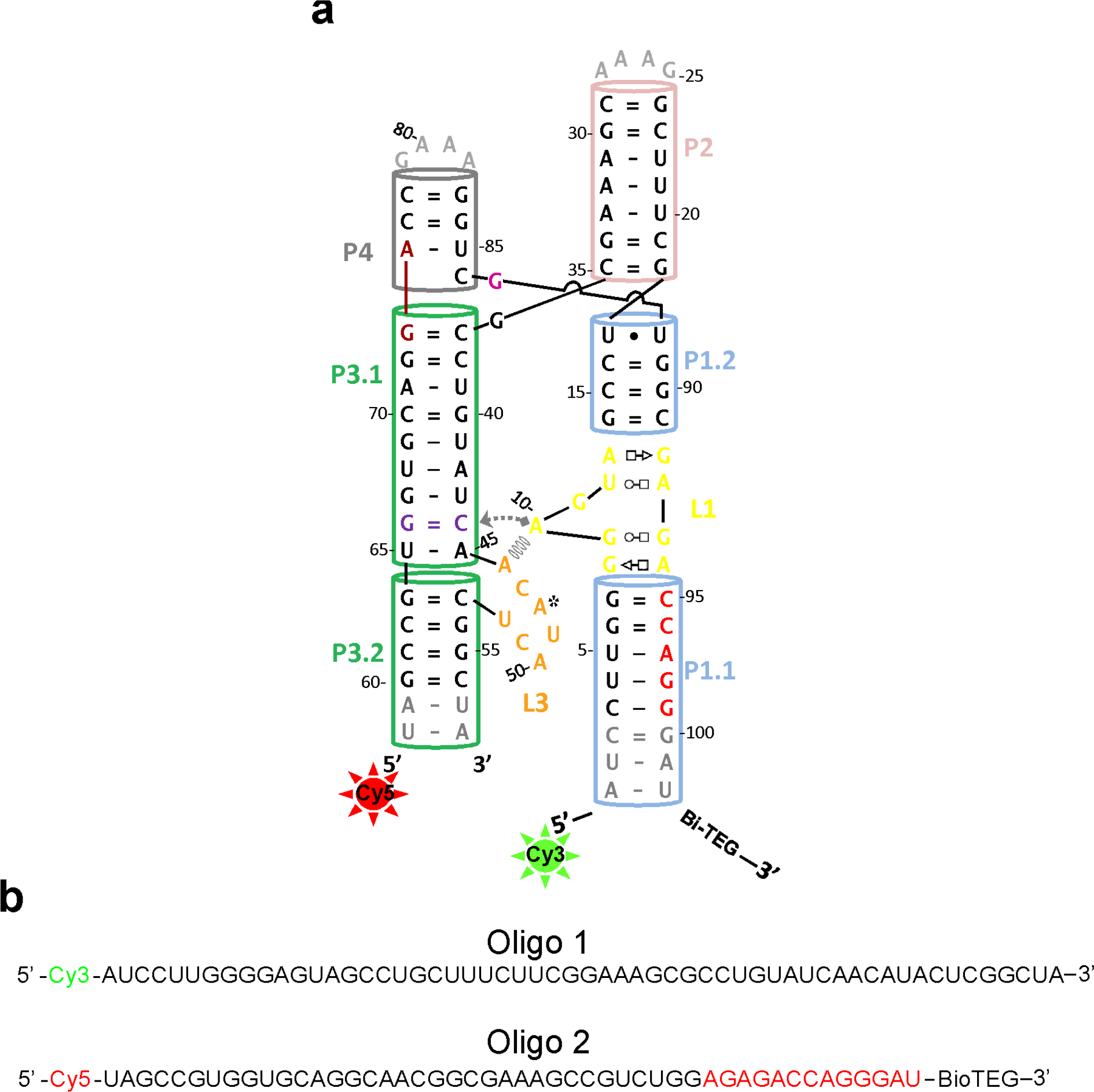
smFRET construct design. (**a**) Sequence and secondary structure of the *Xory yybP-ykoY* riboswitch construct used for smFRET. The construct was made was hybridizing two chemically synthesized RNA oligonucleotides modified at their 5’ and 3’ ends with Cy3, Cy5 fluorophores and Biotin-TEG. (**b**) Sequences of the two oligonucleotides with modifications used for the smFRET construct shown in (a)

**Figure S14.**
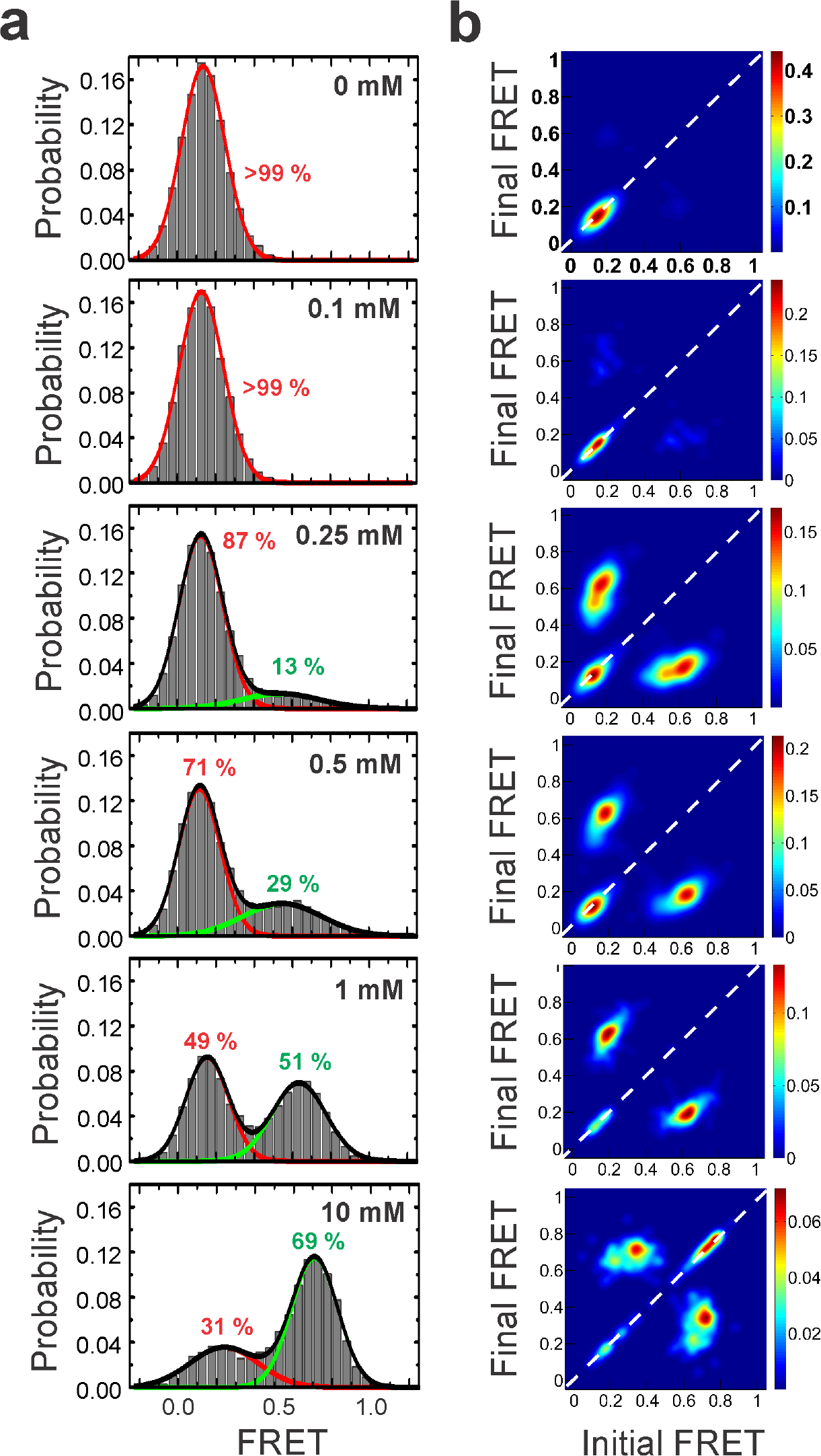
Mg^2+^ titration of the WT *Xory* riboswitch. **(a)** FRET histograms showing the distribution of the two FRET states under various Mg^2+^ concentrations, fit to a sum of Gaussian functions. The Gaussian peaks for the low- and high-FRET states are shown in red and green, respectively, while the cumulative fit is shown in black. The histograms under 0 mM and 0.1 mM contain very low populations of the high-FRET state and were fit to a single Gaussian function. (**b**) TODPs under different Mg^2+^ concentrations showing the fraction of static ‘on-diagonal’ and dynamic ‘off-diagonal’ molecules. The SD populations under the high, 10 mM Mg^2+^ is evident in the TODP.

**Figure S15.**
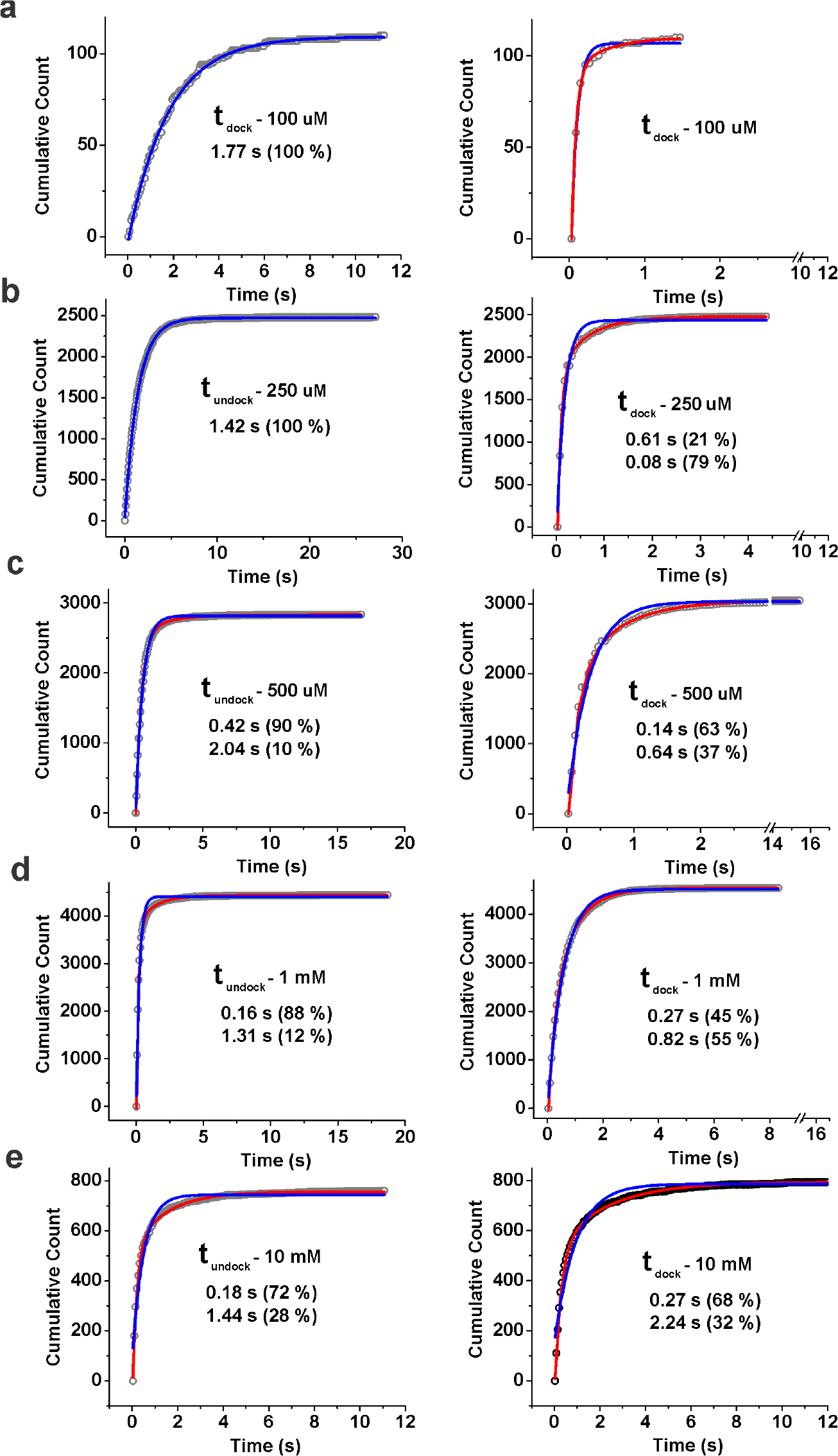
Kinetics of WT *Xory* Mn^2+^ sensing riboswitch. **(a)** Cumulative dwell-time distributions of t_undock_ and t_dock_ in the presence of 0.1 mM MgCl_2_ fit to single (blue) and double-exponential (red) are shown. The life-times and amplitudes of slow and fast components are also shown. In case of double-exponential fits, fit to single exponential function is also shown for comparison. **(b-e)** Same as in (a) but in the presence of 0.25 mM, 0.5 mM, 1 mM and 10 mM MgCl_2_, respectively.

**Figure S16.**
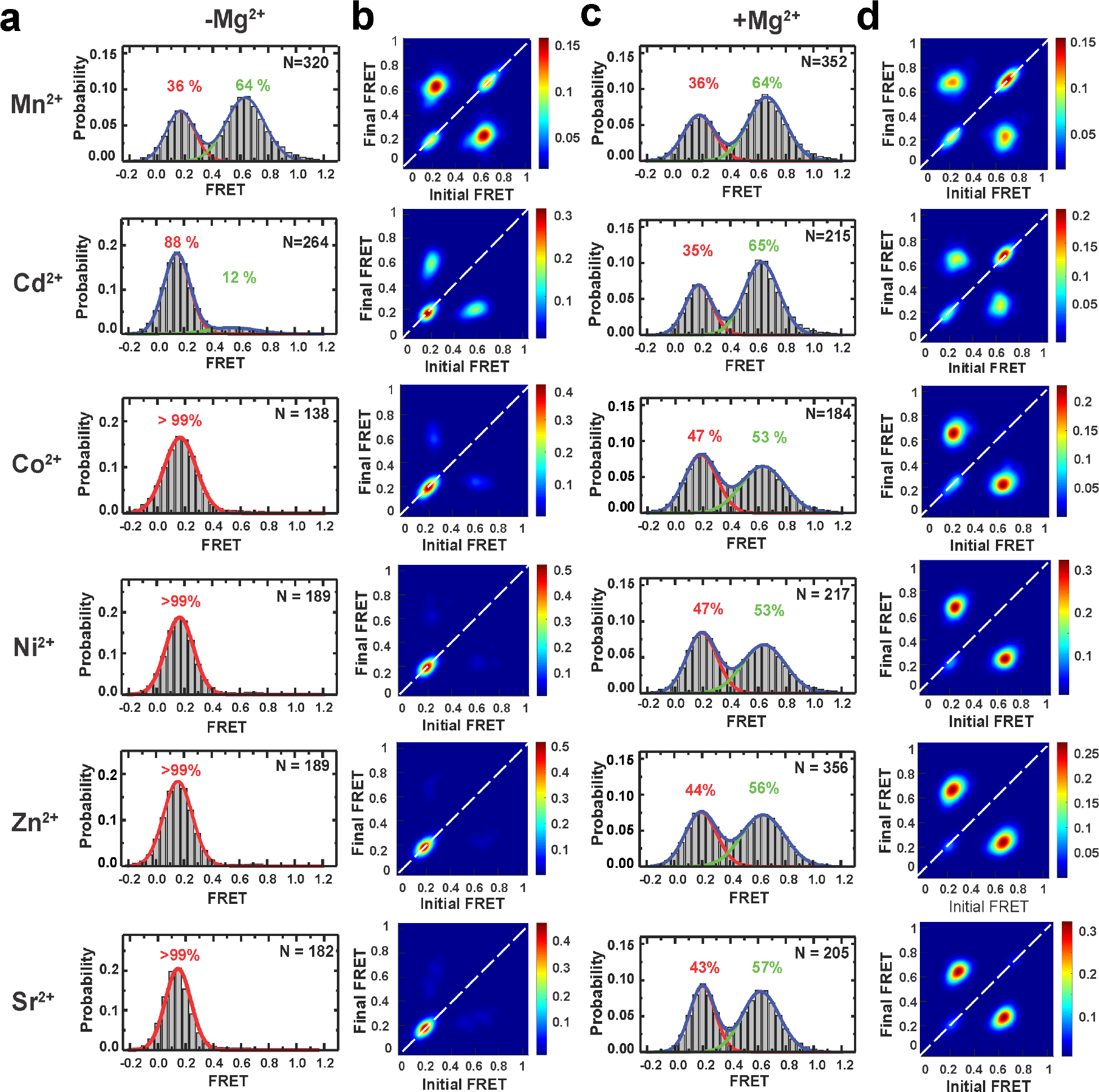
Effect of different metal ions on the WT *Xory* riboswitch. **(a)** FRET histograms showing equilibrium distribution of docked and undocked conformations of the WT riboswitch at 0.1 mM concentration of different transition metal ions alone (i.e., in the absence of 1 mM MgCl_2_). Gaussian fits to the docked, undocked are shown as green and red curves, while the cumulative fit is shown in blue. The number of molecules, N, for each condition analyzed and the % of docked and undocked conformations in each condition are indicated. **(b)** TODPs corresponding to the histograms in (a). **(c)** FRET histograms and **(d)** TODPs at 0.1 mM concentration of different metal ions in the presence of of 1 mM MgCl_2_. For the different metal ions tested, except Mn^2+^ and Cd^2+^, almost all the traces remained in SU conformation, with >95 % in the low-FRET (~0.1) undocked conformation, in the absence of Mg^2+^.

### The *yybP-ykoY* riboswitch can discriminate between similar transition metal ions

The selectivity of the *Xory* riboswitch for Mn^2+^ over Mg^2+^ arises in part from the inner-sphere contact with A48(N7), suggesting that other soft transition metal ions may also be recognized^1^. To test this hypothesis, we probed the effects of different divalent metal ions on the conformation of the riboswitch, at 0.1 mM concentration alone or in the presence of 1 mM Mg^2+^. FRET histograms showed that out of all the different metal ions tested, Cd^2+^ is most effective in promoting docked conformations (**Fig. 5b**, **Supplementary Fig. 16**). In the presence of 1 mM Mg^2+^, addition of 0.1 mM Cd^2+^ resulted in ~65 % of the high-FRET docked conformation, comparable to the docked population upon addition of 0.1 mM Mn^2+^. Examination of individual smFRET traces as well as the TODP showed that this is due to a large fraction of SD traces (**Supplementary Fig. 16**), in agreement with the tight binding of Cd^2+^ to the *yybP-ykoY* riboswitch shown recently^1^. Among the other metals tested, Ni^2+^, Co^2+^, Sr^2+^ or Zn^2+^ had little effect on promoting the folded conformations of the riboswitch under these conditions. Interestingly, in the absence of Mg^2+^, while 0.1 mM Mn^2+^ alone led to the appearance of DD and SD traces with ~62 % docked population (mean FRET 0.67±0.12) (**Fig. 5c**), 0.1 mM of Ni^2+^, Co^2+^, Sr^2+^ or Zn^2+^ did not affect SU traces and Cd^2+^ had only a small effect in promoting DD traces (**Supplementary Fig. 16**). This suggests that, while Mg^2+^ and Mn^2+^ may both bind at M_A,Mg_, Cd^2+^ may be more specific to the M_B,Mn_ site. These results suggest that while the *Xory* riboswitch has some degree of plasticity in recognizing ligands, in a background of Mg^2+^ it preferentially recognizes Mn^2+^ and Cd^2+^ and can effectively discriminate against similar divalent transition metal ions.

**Figure S17.**
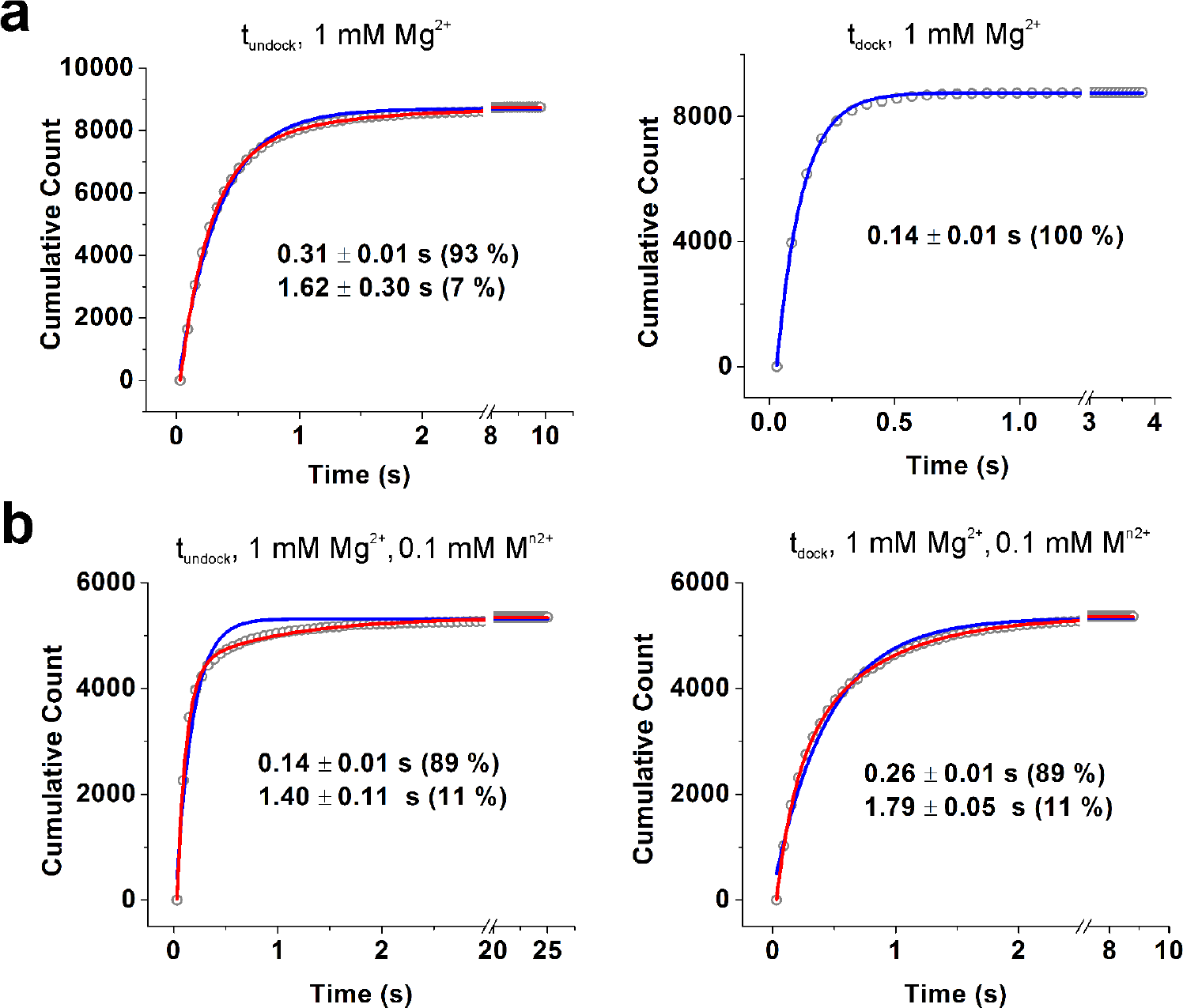
Kinetics of the *Xory* riboswitch A48U mutant. **(a)** Cumulative dwell-time distributions of t_undock_ and t_dock_ in the presence of 1 mM MgCl_2_ fit with single (blue) and double-exponential (red) functions. The life-times and amplitudes of slow and fast components are also shown. In the case of double-exponential fits, fit to single exponential is also shown for comparison. **(b)** Same as in (a) but in the presence of 1 mM MgCl_2_ and 0.1 mM MnCl_2_.

